# The functional neurobiology of negative affective traits across regions, networks, signatures, and a machine learning multiverse

**DOI:** 10.1101/2025.05.15.653674

**Authors:** M. Sicorello, P.J. Gianaros, A.G.C. Wright, P. Bogdan, T.E. Kraynak, S.B. Manuck, C. Schmahl, T.D. Wager

## Abstract

Understanding the neural basis of negative affective traits like neuroticism remains a critical challenge across psychology, neuroscience, and psychiatry. Here, we investigate which level of brain organization—regions, networks, or validated whole-brain machine-learning signatures—best explains negative affective traits in a community sample of 458 adults performing the two most widely used affective fMRI tasks, viewing emotional faces and scenes. Neuroticism could not be predicted from brain activity, with Bayesian evidence against all theory-guided neural measures. However, preregistered whole-brain models successfully decoded vulnerability to stress, a lower-level facet of neuroticism, with results replicating in a hold-out sample. The neural stress vulnerability pattern demonstrated good psychometric properties and indicated that negative affective traits are best represented by distributed whole-brain patterns related to domain-general stimulation rather than localized activity. Together with results from a comprehensive multiverse analysis across 14 traits and 1,176 models— available for exploration in an online app—the findings speak against simplistic neurobiological theories of negative affective traits, highlight a striking gap between predicting individual differences (*r*<.35) and within-person emotional states (*r*=.88), and underscore the importance of aligning psychological constructs with neural measures at the appropriate level of granularity.

---

People differ in their tendency to experience negative emotions, which is reflected in all contemporary models of human personality and psychopathology under various labels like “negative affectivity”, “neuroticism”, “negative affect”, “affective instability” (Brandt & Mueller, 2022; DeYoung et al., 2022; Eysenck, 1967; Gore & Widiger, 2018; Shackman et al., 2016; Wright & Simms, 2015). As a cornerstone of personality models ((McCrae & Costa Jr., 2008) and novel dimensional taxonomies for mental disorders ((Kotov et al., 2017); (Cuthbert, 2014; Kozak & Cuthbert, 2016), identifying the biological basis of negative affectivity is crucial for refining theoretical models by providing physical counterparts to latent psychological trait constructs. This serves the discovery of novel targets for personalized biologically oriented interventions such as psychotropic medication, neurofeedback and (non-)invasive brain stimulation (DeYoung et al., 2022). Still, despite extensive theoretical contributions and empirical research, the neurobiology underlying negative affectivity remains largely unclear.

There have been three levels for neural targets of negative affectivity, which differ in complexity and spatial scale: (1) Single brain regions, (2) large-scale networks based on resting state-data, and (3) validated whole-brain neural signatures (Figure 1a). For the single region approach, the most common targets are the amygdala, the anterior insula, and the dorsal anterior cingulate cortex (dACC), as they are activated by tasks involving negative stimuli in both human and animal models (Shackman et al., 2016). Of these regions, the amygdala has received the most attention (Eysenck, 1967; Shackman et al., 2016). At the second, network level of analysis, these three regions also represent nodes of the so-called “salience network”, which is similarly often taken to infer negative affectivity (Seeley, 2019). Concerning the third neural level, validated whole-brain neural signatures represent multi-system machine learning-based models, which are trained to be strongly associated with self-reported affective states during experiments, and typically include regions across multiple networks (P. A. Kragel et al., 2018).

**Figure 1.**
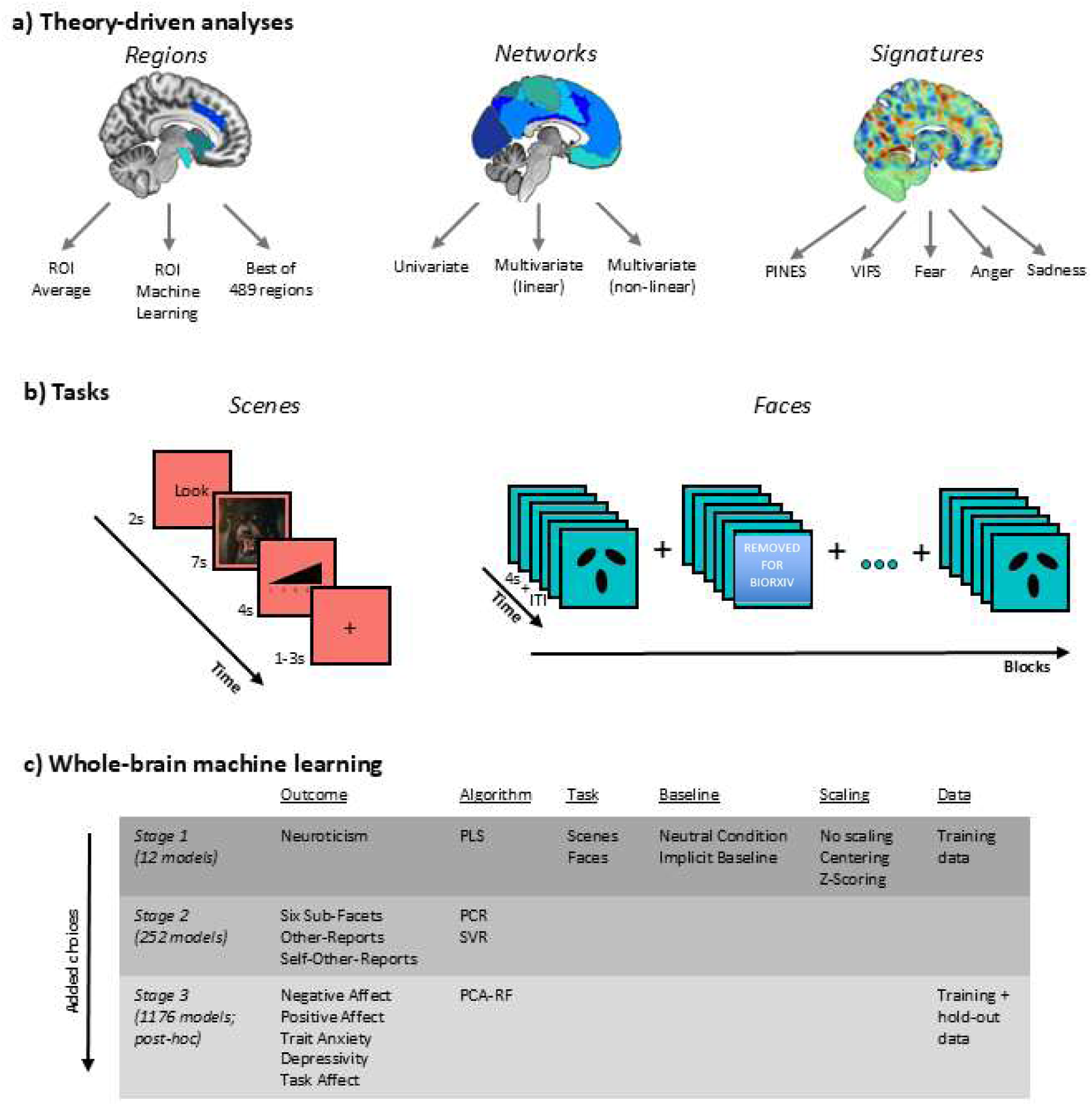
Schematic illustration of the study design. (a) Theory-driven analyses for regions, networks, and signatures. The three ROIs (amygdala, anterior insula, dACC) are extracted from different sagittal slices and projected on the same slice for visual simplicity. ROI = region of interest, PINES = Picture Induced Negative Emotion Signature, VIFS = Visually Induced Fear Signature. (b) Structure of scenes and faces task. (c) Whole-brain machine learning models across three sequential analytic stages (first two preregistered). Each stage adds modeling choices to the previous stages. PLS = Partial Least Squares Regression, PCR = Principal Component Regression, SVR = Support Vector Regression, PCA-RF = Principal Component Analysis with Random Forest Regression.

Process-oriented research has convincingly demonstrated the shortcomings of single brain regions and networks for the inference of negative affective *states* (i.e., observing the same individuals under different conditions). Besides findings that regions and networks are domain-general and therefore not specific to negative affective states (i.e., also respond to non-emotional or positive stimuli; (Cunningham & Brosch, 2012; Lindquist et al., 2016; Seeley, 2019; Uddin, 2015), several well–powered studies have shown they only moderately track self-reports of negative affective states at best (Chang et al., 2015; P. A. Kragel et al., 2018; Krishnan et al., 2016; Zhou et al., 2021). Specifically for the amygdala, even evidence for its involvement in fear learning and extinction is inconsistent based on human fMRI data (Fullana et al., 2018; Taschereau-Dumouchel et al., 2019; Visser et al., 2021) and deep-brain stimulation does not clearly invoke conscious states of fear (Inman et al., 2020). In contrast, neural signatures can reach very large cross-validated correlations with self-reported negative affective states (*r*=.92; (Chang et al., 2015) and self-reported fear (*r*=.82; (Taschereau-Dumouchel et al., 2019; Zhou et al., 2021), which generalize to clinical datasets (Sicorello et al., 2021), all while maintaining specificity from other negative mental events like physical pain (Čeko et al., 2022). Taken together, while some neural signatures effectively track negative affective *states*, single regions and networks do not, making the interpretation of these neural targets more ambiguous.

The literature on associations of specific regions and networks with affective *traits* is less clear. On the one hand, there is a large literature reporting a convergence between negative affectivity and the brain regions concerned with salience processing (Mitchell & Kumari, 2016; Ormel, Bastiaansen, et al., 2013; Shackman et al., 2016). Moreover, several meta-analyses found significant associations between affect-related mental disorders for the amygdala, anterior insula and the dACC, based on structure, functional connectivity, and responsivity to negative stimuli (Goodkind et al., 2015; Schulze et al., 2019; Sha et al., 2019). On the other hand, the transdiagnostic effects found in these case-control studies could be explained by a large number of confounding factors, and thus may not be specific or even directly related to negative affectivity. In addition, meta-analyses on dimensional negative affectivity traits do not converge on a single set of theory-driven regions or networks in terms of neural responsivity to negative stimuli, structure, or functional connectivity (Y.-W. Chen & Canli, 2022; Lin et al., 2023; Liu et al., 2021; Mincic, 2015; Sicorello & Schmahl, 2021; Xu et al., 2019).

A major shortcoming of the literature is the combination of very small sample sizes in single studies and very low test-retest reliability for region-based neural measures (Elliott et al., 2020; Noble et al., 2019; Szucs & Ioannidis, 2017; Zugman et al., 2023). Large-scale efforts have found only very small associations between individual differences in dimensional psychopathology and brain structure or resting connectivity (Marek et al., 2022; Schulz et al., 2024)*R*²=0.01-.03; (Marek et al., 2022; Schulz et al., 2024). Nevertheless, psychological theory and evidence from daily life indicate negative affectivity largely manifests in an increased reactivity to stressful or threatening environments (Bolger & Schilling, 1991; Bolger & Zuckerman, 1995; Ringwald et al., 2024; Shackman et al., 2016). Hence, it is possible that neural correlates of negative affectivity can mainly be measured in appropriate task contexts (Brandt & Mueller, 2022). Still, a study on over 500 young adults did not find an association between neuroticism and amygdala responsivity when viewing negative faces (Silverman et al., 2019), supporting an earlier meta-analysis (Servaas et al., 2013). In contrast to single regions or networks, whole-brain neural signatures for negative affect not only correlate much more strongly with momentary self-reports, but also have much higher between-run and test-retest reliability (Han et al., 2022; P. Kragel et al., 2020; Sicorello et al., 2021). Despite these advantages, previous work testing relationships between individual differences and responses in affect-related neural signatures has not yet proven successful (Sicorello et al., 2021). However, a dimensional approach covering the full spectrum of negative affectivity traits is still missing. As psychological traits have a hierarchical structure, from broad domains to specific facets, it is crucial to determine which trait level corresponds to neurobiological processes in common tasks (Brandt & Mueller, 2022).

Here, we present a comprehensive preregistered in-depth investigation of the neural correlates of negative affectivity across different trait constructs, neural levels, and the two most widely used affective fMRI tasks: viewing pictures of negative faces and scenes.

First, we present theory-driven associations between neuroticism and neural regions, networks, and signatures, testing widespread ideas from the current and past literature (Figure 1a). We used neuroticism as a well-studied exemplar of negative affectivity as the main outcome due to its strong theoretical basis, its large associations with other negative affectivity constructs, its relevance for mental and physical health, and its hierarchical subfacets which allow more fine-grained approaches (Cuijpers et al., 2010; Gore & Widiger, 2018; Ormel, Bastiaansen, et al., 2013; Ormel, Jeronimus, et al., 2013). The region approach includes both average signal and multivariate patterns within the amygdala, anterior insula, dACC, as well as data-derived best predictive regions. The network approach covers the average effect of single networks, but also linear and non-linear network interactions, based on neurobiological theories of emotion (Lindquist & Barrett, 2012). The neural signatures approach covers well-validated neural signatures for negative affect, fear, and discrete negative emotions, which are heavily understudied for individual difference research (Chang et al., 2015; P. A. Kragel & LaBar, 2015; Zhou et al., 2021).

Secondly, we present data-driven machine learning analyses based on whole-brain data to derive a multivariate signature for negative affectivity and validate it in a hold-out sample. Traits like neuroticism are higher-order constructs, which possibly do not match the abstraction level of specific task-based neural measures (Brandt & Mueller, 2022). Therefore, additionally to neuroticism, we also tested the predictability of the six lower order facets of neuroticism (depression, anxiety, stress vulnerability, anger, impulsiveness, self-consciousness). The neural signature for the construct with the highest accuracy was further assessed for its psychometric properties (reliability and validity), psychological and neurobiological interpretability, and its relation to theory-driven neural measures. These steps are crucial to provide a viable neural marker indicate of negative affective constructs.

Lastly, we present extensive exploratory multiverse analyses (Simonsohn et al., 2020) on the predictive accuracy of ≈1200 models across a larger set of 14 affective trait constructs as well as other design and data analytic choices, which we made available via an interactive online app to allow the free exploration of design factors influences predictive accuracy.

In the preregistration, we hypothesized that a whole-brain predictive pattern for neuroticism would outperform region-, network-, and signature-based models. Given the proposed importance of aligning fMRI tasks with psychological constructs, we expected this pattern to (1) show greater accuracy for the scenes task than the faces task, as the latter also taps into emotion recognition; (2) correlate more strongly with negative affect than with positive affect or extraversion; and (3) primarily reflect withdrawal-related facets of neuroticism, including depression, anxiety, and stress vulnerability.

The timestamped preregistration (October 2020) of aims, hypotheses, and statistical analyses as well as data, analysis code, and trained model patterns can be found on: https://github.com/MaurizioSicorello/NeuroSquare_repo.

## Methods

### Participants

The data comprised baseline assessments from up to 424 participants from the Adult Health and Behavior project (phase 2; AHAB-2) and the Pittsburgh Imaging Project (PIP). Therefore, this method section partially overlaps with details from previous publications (especially (Gianaros et al., 2020)], which used identical MRI preprocessing). All 424 participants provided data for the faces task, while only a subset of 338 participants provided data for the scenes task. Sample characteristics are provided in table 1. All participants provided informed consent. The University of Pittsburgh Human Research Protection Office granted approval for AHAB-2 (Protocol ID: 07040037) and PIP (07110287), as well as their aggregation to create a common data registry (19030174).

**Table 1.** Sample Characteristics.

|  | faces (N = 424) | scenes (N = 338) |
| --- | --- | --- |
| Women (%) | 52.1 | 48.8 |
| Age (years) | 42.8 ± 7.4 | 41.3 ± 7.1 |
| Race (%) |  |  |
| Caucasian/White | 83.5 | 77.7 |
| African-American | 15.1 | 19.0 |
| Multiracial/ethnic | 1.4 | 1.2 |
| School years completed | 17 ± 2.8 | 16.6 ± 3 |

### Psychological measures

Neuroticism and extraversion were measured with self reports from the full the NEO-PI-R (Costa Jr. & McCrae, 2008), and “other-reports” from up to two close information in AHAB-2 using the NEO-FFI (Costa & McCrae, 1992); spouses/partners, first-degree relatives, close friends and co-workers). Positive and negative affect were measured using the Positive and Negative Affect Schedule (Crawford & Henry, 2004). Trait Anxiety was measured with the Trait-State Anxiety Inventory (Spielberger, 1989). Dimensional depression severity was measured with the Beck Depression Inventory (Beck et al., 1996). Trait-like responsivity to negative stimuli was measured as the person-wise average difference in self-reported momentary affect, based on a single-item 5-point Likert scale (see task for further information). Descriptive statistics and correlations are shown in Table S1. The main outcome, neuroticism, has an estimated internal consistency of *r* = .93 and test-retest reliability over three months of *r* = .79 (Costa & McCrae, 1992). Internal consistency of subfacets is slightly lower around *r* = .80.

### Experimental tasks

Schematics of task structures are depicted in Figure 1b for both tasks.

#### Scenes task

Subgroups of AHAB-2 and PIP participants completed an affective processing and responding task, which has been detailed previously (Gianaros et al., 2014). Participants saw 30 unpleasant and 15 neutral IAPS images (Lang et al., 2008). Participants were first trained and then instructed to (i) ‘Look’ and attend to images or (ii) ‘Decrease’ and change their thinking about the image to feel less negative. Trials were comprised of a 2s cue (‘Look’ or ‘Decrease’), followed by a 7s IAPS image presentation. After image viewing, participants rated their emotional state (‘How negative do you feel?’) on a 5-point Likert-type scale in a 4s rating period (1 = neutral, 5 = strongly negative). A variable (1–3s) rest period followed each rating period. The entire task duration was 11 min and 16s (15 ‘Look neutral’ trials; 15 ‘Look negative’ trials; 15 ‘Decrease negative’ trials). Images were presented such that no more than two identical trial types (‘Look negative’ or ‘Decrease negative’) were consecutive and no more than four unpleasant images were consecutive. Given the focus of this report (and for comparability to the facial expression tasks involving the passive viewing of affective stimuli), only ‘Look neutral’ and ‘Look negative’ trials were analyzed. Across AHAB-2 and PIP, unpleasant images for ‘Look negative’ trials overlapped by 87% (13/15 shared images) and neutral images by 13% (2/15 shared images). E-Prime software (Psychology Software Tools, Sharpsburg, PA) was used to present stimuli and record behavioral responses in the facial expression and IAPS tasks (for a list of image stimuli, see Gianaros et al., 2020).

#### Faces task

In this task, participants completed four blocks of a facial expression-matching-to-sample condition, which was interleaved with five blocks of a shape-matching (sensorimotor) control condition. In the facial expression matching condition, participants saw three same-sex faces in an array for each trial, all expressing either fear or anger. Participants chose one of the two faces at bottom that was identical to a center target at top. Each block consisted of six trials (three fear, three anger; three all-male, three all-female), and each trial lasted for 4s (1.5, 3.5 and 5.5s variable inter-trial interval; ITI). All images were in black and white and were drawn from the Pictures of Facial Affect (PFA) stimulus set (Ekman & Friesen, 1976). In the control condition, participants also matched-to-sample but instead used images of circles, vertical ellipses and horizontal ellipses. Each trio of shapes was shown for 4s with a 2s ITI. The total task length was 6 min and 36 seconds, including 6s for initial magnetic equilibration. Similar tasks have been used frequently in the study of emotion processing and psychopathology, including the large-scale UK biobank and ABCD studies (Casey et al., 2018; Miller et al., 2016).

### MRI data acquisition and analysis

Functional blood-oxygen-level-dependent (BOLD) images from AHAB-2 and PIP participants were collected on the same 3 Tesla Trio TIM whole-body scanner (Siemens, Erlangen, Germany), equipped with a 12-channel head coil. Further details on data acquisition and pre-processing can be found in the supplements. Univariate general linear models (GLMs) were estimated to compute condition or event contrast maps that were later used for prediction analysis. Stimulus timeseries were convolved with the default hemodynamic response function in SPM12. For the facial expression matching task, these regressors modeled blocks of face- and shape-matching conditions using boxcar functions. For the IAPS task, these regressors modeled events of the trial (i.e. cue, picture, rating period, rest). Each GLM additionally included six motion regressors of no interest and a high-pass temporal filter (128s) to correct for low frequency drift. Group mean contrasts were estimated using restricted maximum likelihood, as implemented in the RobustWLS toolbox, v4.0 (Diedrichsen & Shadmehr, 2005). These included the ‘Faces vs Shapes’ contrast for the facial expression-matching tasks and ‘Look negative vs Look neutral’ contrast for the IAPS task.

### Data Exclusion

One participant in the faces task was a multivariate outlier according to Bonferroni-Holm-corrected Mahalanobis distance and therefore excluded from analyses on that task. The sample size reported throughout this manuscript reflects the effective (i.e., final) sample size.

### Statistical Analysis

#### Power analysis of hold-out samples

A hold-out sample of N = 102 for the scenes task and of N = 101 for the faces task was drawn for the evaluation of machine learning models. Training and hold-out samples were stratified for neuroticism using the caret package in R (v4.3.2) to make the two subsamples comparable. The discrepancy of one participant was due to the different number of participants for the two tasks and rounding in the stratification procedure. This sample has a power of .90 to detect a true correlation between pattern and neuroticism of r ≅ .30 in a one-tailed test, which is at the 75% percentile of questionnaire-behavior correlations in individual difference research (Gignac & Szodorai, 2016).

#### Theory-Driven Approaches

For the theory-driven approaches, neural activity was assessed in the contrasts for negative vs neutral scenes and negative faces vs shapes. An overview of models is illustrated in Figure 1a.

##### Regions

Multiple atlases were used to parcellate the brain into 489 anatomic regions taken from the canlab toolbox (Geuter et al., 2020; Wager, 2024). From these, masks for the amygdala, anterior insula, and dACC were built. We conducted three region-based tests, which vary in complexity. First, correlations between average signal in these regions and neuroticism scores was calculated on the full data, separately for the two hemispheres and Bonferroni-Holm corrected for six tests. Secondly, we predicted neuroticism scores from voxel-wise activity in each region in the training sample using cross-validated partial least squares regression and evaluated performance in the hold-out sample. We used the same procedure as described in the machine learning section below. This was done to account for the fact that using only the average signal of these regions might be too coarse. Thirdly, we extended our analysis to regions beyond the three main regions. We split the sample in half, stratified for neuroticism, picked the region with the strongest correlation in the first half and tested its correlation in the second half.

##### Networks

A canonical network parcellation was used to extract the average signal in seven resting state networks (Thomas Yeo et al., 2011): Limbic, Ventral Attention (Salience Network), Dorsal Attention, Fronto-Parietal, Default-Mode, Visual, and Somatomotor. Again, we used three different approaches, which differ in complexity. First, we correlated this signal with neuroticism in the full sample to test the simple hypothesis that one of these networks predominantly underlies negative affectivity, correcting for seven tests using the Bonferroni-Holm procedure (the preregistration stated an incorrect correction for eight tests). Then, we predicted neuroticism from average activity in all seven networks using multiple regression to test whether neuroticism is represented by a linear combination of network activities. Last, we repeated this predictive model with random forest regression to allow for non-linear effects and interactions between networks. For the latter model, performance was evaluated using the out-of-bag samples (a form of cross-validation) and a permutation test with 1000 random permutations of neuroticism scores.

##### Signatures

We calculated the neural expression of five well-validated neural signatures for negative affect (PINES; (Chang et al., 2015; P. A. Kragel & LaBar, 2015; Zhou et al., 2021)), fear (VIFS;(Chang et al., 2015; P. A. Kragel & LaBar, 2015; Zhou et al., 2021)), and three discrete negative emotions (Chang et al., 2015; P. A. Kragel & LaBar, 2015; Zhou et al., 2021) by taking the dot product of model weights and person-wise functional images. The VIFS was published after our preregistration and therefore added post-hoc. Moreover, we preregistered an analysis combining the three discrete emotions into one factor score for “neural negative affect”, before assessing them separately, but their intercorrelations were too low to make this a sensible approach (all r < .12). A Bonferroni-Holm correction for three tests was preregistered for the discrete emotion patterns, but a correction for five tests is also sensible.

#### Data-Driven Machine Learning Models

The general preregistered strategy was to (1) split the data into a training and a hold-out sample, (2) predict neuroticism from different models in the training sample using nested cross-validation, (3) retrain the best model on the entire training dataset, and (4) assess performance in the hold-out sample. An overview of all modeling choices can be found in Figure 1c. Machine learning analyses were performed using the CanlabCore Toolbox (Wager, 2024)

In step 1, the hold-out sample was set aside with the procedure described in the power analysis section.

In step 2, we predicted neuroticism in the training sample using Partial Least Squares regression from a small set of 12 “first stage” models (Figure 1c) with different combinations of task choice (scenes vs faces), baseline choice (neutral scenes or shapes vs implicit baseline), and image-wise scaling choice (z-scoring vs centering vs no scaling). All whole-brain machine learning models were conducted after applying a gray matter mask and z-standardizing voxel-wise values on the sample mean and standard deviation of that voxel. As other studies will have different voxel-wise statistics, predictions from our model will have an offset and can only serve as a correlate of trait measures, rather than predicting exact numeric values. Repeated nested cross-validation (2×5×5) was used to calculate cross-validated correlations between pattern predictions and outcome scores, with five inner folds for hyperparameter tuning, five outer folds for model evaluation, and two repeats for stability. Pearson’s correlations were used as a performance metric, as the aim was to train a pattern for scale-independent correlates of negative affectivity, which can be used across different studies with different scanning parameters or different questionnaires. We preregistered that if the best-fitting model reaches p < .05 (one-tailed) in a permutation test, the model is accepted and will be validated in the hold-out sample, again with a one-tailed p < .05 and supplemented with a Bayes factor with default low-informative half-cauchy priors for one-sided tests from the Bayes factor package in R (Morey & Rouder, 2018). One-tailed tests were used, as only positive correlations indicate successful prediction. If this criterion was not fulfilled, we preregistered a plan to move on to a larger set of 252 “second stage” models (Figure 1c), comprising several choices of outcome (neuroticism and its six facets) and algorithms (Partial Least Squares [PLS],Principal Component Regression [PCR], and Support Vector Regression [SVR]). We also preregistered the inspection of self- and other-based ratings, as well as their combination. For further information on hyper-parameter estimation and model evaluation see the supplements. The machine learning procedure was tested with a simulated dataset of *N* = 100 and 20 predictive features, implemented as a “test mode” in the main script.

#### Exploratory Multiverse Analysis

In a post-hoc multiverse analysis, we calculated a larger set of 1176 “third stage” models (Figure 1c) to assess the effect of different negative affectivity traits and modeling choices on predictive accuracy. Additional outcomes included negative affect, positive affect, STAI scores, BDI scores, other-reports for neuroticism, and task-based affective self-ratings collected during the IAPS task (person-wise averaged for the negative minus neutral condition). Other modeling choices included using the full data (training plus hold-out) to assess the effects of sample size and using random forest regression as an algorithm which accommodates nonlinear and interaction effects. For random forest regression, principal component analysis was used to reduce the feature space to the number of participants in the training data.

The impact of design choices was assessed by predicting brain-outcome-correlations from design factors using random forest regression with 1000 bagged trees (2×5=10 correlations per model). The relevance of single design predictors were quantified with permutation-based variable importance. These analyses were conducted with the cforest package in R. Moreover, we performed a variance decomposition to quantify which design factor is associated with the largest differences in cross-validated correlations. These analyses were only performed with descriptive rather than inferential statistics, as the latter might be incorrect due to complex dependencies in the data (Bates et al., 2024).

As this number of models was already very large and computationally intensive, we omitted the additional preregistered choices of image-type (beta-map versus t-map), outcome modelling (sum scores versus factor scores), and the neuroticism spectra withdrawal and volatility in all analyses, as we believed these choices to have little impact on performance (beta-images and t-images are highly correlated, all information of neuroticism spectra is contained in the more fine-grained facets, and factor scores, in contrast to latently modeled variables, also contain measurement error and are extremely highly correlated with sum scores; (G. Chen et al., 2017; Widaman & Revelle, 2023).

## Results

### Theory-Driven Approaches: Regions, Networks and Signatures

Overall, there were no clear associations between neuroticism and any theory-driven brain measure across regions, resting-state networks, and signatures (Figure 2). Most confidence intervals ruled out correlations smaller than |r| = .20 and even uncorrected Bayes factors mostly favored the null hypothesis with values above 10. For precise point estimates, confidence intervals, and Bayes factors see Table S2.

**Figure 2.**
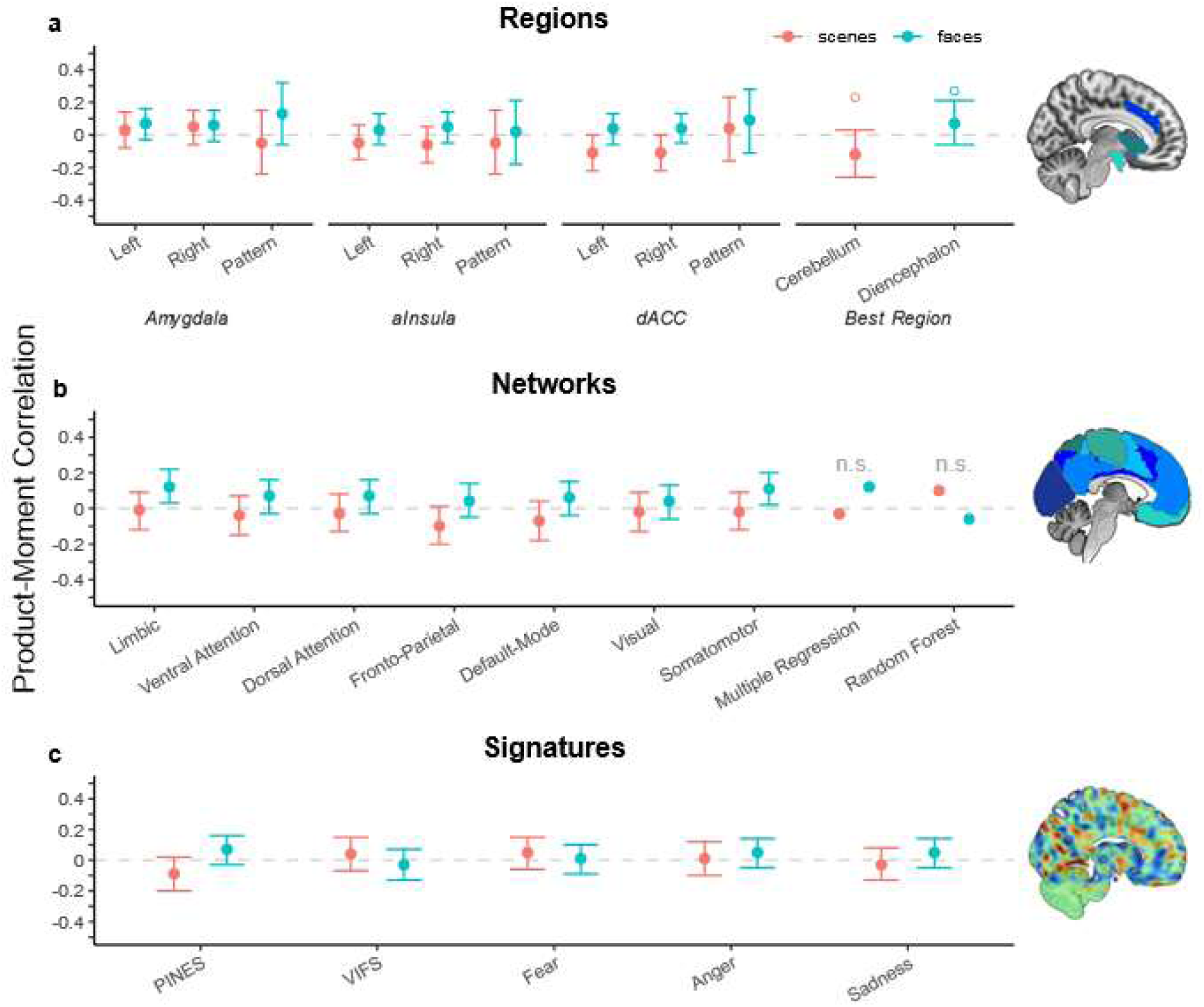
Correlations between neuroticism and brain measures across two fMRI tasks and three neural levels: Regions, networks, and validated multivariate neural signatures. For region sub-models, “pattern” refers to machine learning models limited to these regions. Hollow circles for the best region approach indicate the point estimates from the split-half selection sample.

For regions, amygdala, anterior insula, dACC and a whole-brain discovery procedure were used. Average bilateral activation in the dACC during negative versus neutral scenes was closest to a conventional uncorrected significance threshold (left: *p* = .051; right = .041), but did not survive the preregistered correction for six tests and had signs contrary to theoretical expectations (i.e., lower dACC reactivity was correlated with higher neuroticism). In local region-based multivariate pattern analysis, predictive accuracy was not above chance (all *p* > .195) and 49-51% of voxels had a positive sign, indicating that the direction of the association between activity and neuroticism was not consistent across voxels. The best-single-region approach picked a cerebellar region for the scenes task and a region in the diencephalon for the faces task, but both effects were not significant in the test-half of the sample and the cerebellar effect even changed directions (Table S2).

For networks, only the limbic (*r* = .12, *p* = .011) and somatomotor network (*r* = .13, *p* = .020) crossed an uncorrected threshold in the faces task, but again did not survive a correction for the number of tested networks. Notably, we observed that the activity of all seven networks was very highly correlated on a between-person level (scenes: average *r* = .68, range = [.48- .84]; faces: average *r* = .67, range = [.48-.88]). Hence, a person with higher activity in one network also tended to have higher activity in other networks, compared to other people.

For neural signatures–the most reliable and psychologically valid brain measures of within-person negative affective responses–no tests of associations with neuroticism were statistically significant (all *p* > .09 and |r| < .10). Only the PINES signature for negative affect had Bayes factors lower than 15:1 in favor of the null, with a value of 9.5:1 for faces and 5.5:1 for scenes. However, correlation between neuroticism and PINES responses to IAPS images was negative (*r* = -.09, *p* = .092), indicating smaller reactivity in participants with higher neuroticism.

In sum—using Bayesian inference, confidence intervals, and cross-validation—we found substantial evidence against meaningful theory-driven associations between neuroticism and common brain measures on a neural level of regions, networks, and well-validated affective signatures.

### Data-Driven Approaches: Whole-Brain Machine Learning Prediction

#### Model identification and performance

None of the first stage whole-brain machine learning models (Figure 1c) had positive cross-validated correlations with neuroticism in the training sample, making a permutation test superfluous (best model: *r* = .00). Of the second stage models (Figure 1c), the best model predicted the neuroticism facet *vulnerability to stress* at *r* = .21 (*p* = .009; one-sided) from the scenes task and the contrast [negative – neutral] using SVR and image-wise centering. In the hold-out sample, this correlation was still statistically significant (one-sided): *r*(100) = .19, *p* = .028, *ΒF*_10_ = 2.53, 95% CI [-.00, .37]. The Bayes factor was 2.5:1 in favor of the alternative hypothesis, and crossed a threshold of 3 only when a more narrow default prior is assumed, corresponding to the expectation of smaller effect sizes (*ΒF*_10_ = 3.09). Notably, the detected effect size corresponds to the median effect size of general individual difference research (Gignac & Szodorai, 2016).

For the vulnerability facet, cross-validated accuracy was considerably higher in the scenes than the faces task, but there was no consistent general superiority of all withdrawal-related facets (Figure 3a). Interestingly, other-reports had larger accuracies than self- or combined reports (Figure 3b).

**Figure 3.**
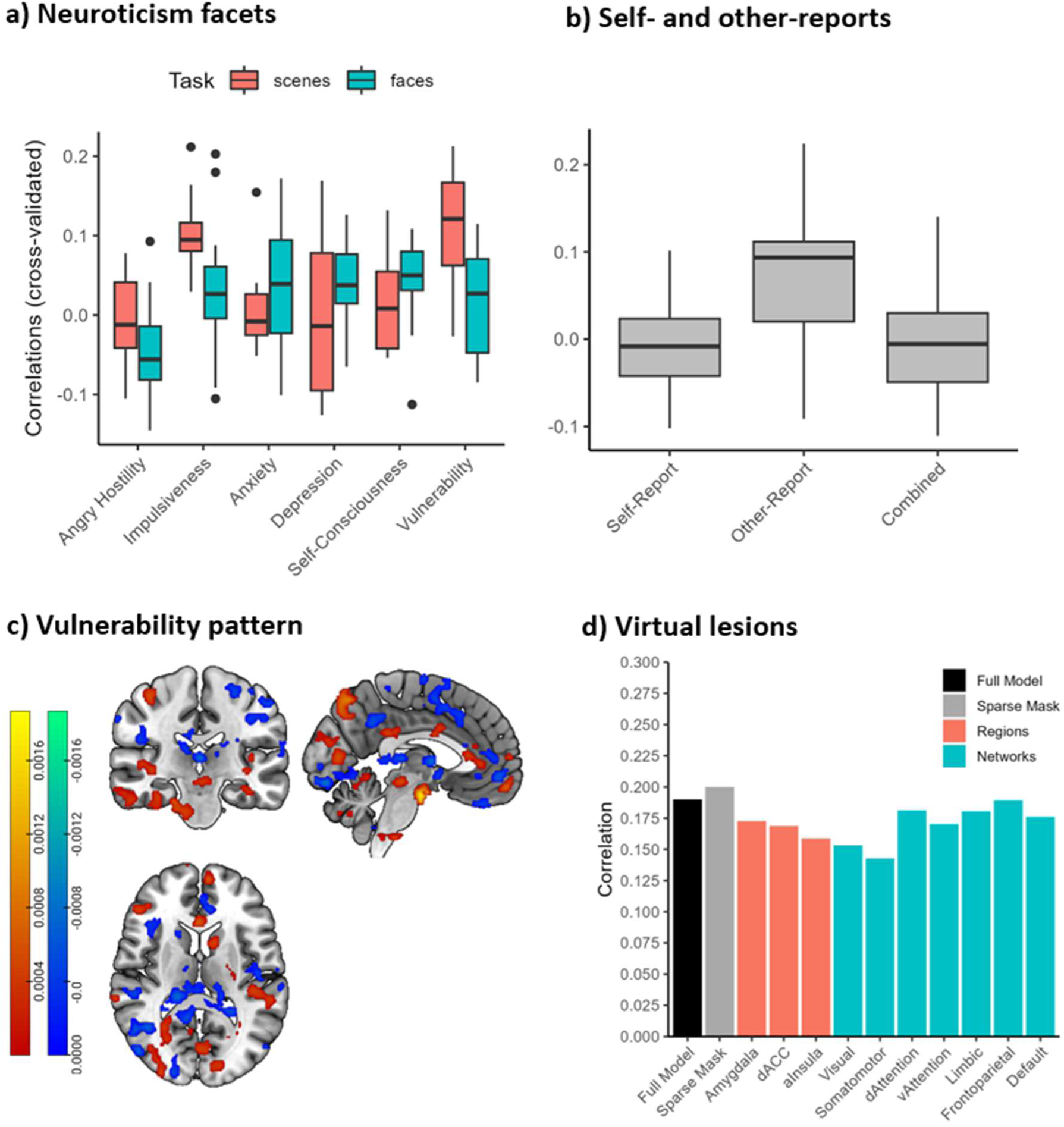
Cross-validated accuracy in the training data as a function of (a) neuroticism facet and task and (b) self-reports vs reports of close others. Includes only preregistered models. (c) Significant regression weights for the stress vulnerability pattern after correction at α = .05 and minimum cluster size of 50 (positive weights in red, negative in blue). (d) Correlations between brain pattern and hold-out data (N=102) after removing single regions and networks. “Sparse mask” represents the performance of the full model after applying a more conservative gray matter mask, which decreases the occurrence of artifactual voxels but also has a higher probability to miss relevant voxels.

Taken together, brain responses to negative scenes could predict the neuroticism facet vulnerability to stress, replicating in the hold-out set. Therefore, person-wise activity of this neural pattern might serve as a neural marker in clinical and personality research. In contrast, prediction of the broad neuroticism domain was not possible, highlighting the importance of focusing on more specific affective traits at lower granularity levels. This neural pattern of stress vulnerability was further assessed for psychometric properties, neurobiological interpretability, and in relation to the theory-driven measures of the section above.

#### Psychometrics

A useful personalized biological measure must be both reliable and valid. The split-half reliability of the person-specific stress vulnerability pattern expression was *r* = .83 (Spearman-Brown corrected) and therefore sufficiently high for psychometric purposes, especially compared to the split-half reliability of the amygdala responses (*r* = .29). We assessed the construct validity by correlating cross-validated pattern predictions in the full sample with trait negative affect (convergent validity) as well as trait positive affect and extraversion (discriminant validity) using leave-one-out cross-validation. For reference, the correlation between pattern predicted and actual stress vulnerability scores were at *r* = .25 using this procedure. Predictions were positively correlated with trait negative affect (*r* = .13), but not trait positive affect (*r* = .01) or extraversion (*r* = .00), indicating a degree of convergent and discriminant validity. We further tested the similarity of the stress vulnerability pattern with 50 topic maps related to psychological processes (e.g., working memory, emotion, executive control) derived from 11,406 previous fMRI studies using Neurosynth (Yarkoni et al., 2011). Neurosynth decoding indicated that brain activation implied by our pattern is most similar with fMRI studies researching the broad psychological topic “stimulation” (*r* = .26).

#### Neurobiological Assessment

The spatial correlation between the vulnerability pattern and the five theory-driven affective neural signatures were all below |r| = .02, which is in line with our null findings for the affective signatures in the theory-based section. No voxel-wise regression weights of the vulnerability-pattern passed FDR correction at q = .05. To still give some intuition about the location of the most strongly contributing voxels, we applied a more lenient thresholding procedure used in a previous study on this dataset at α = .05 and a minimum cluster-size of 50 contiguous voxels (Gianaros et al., 2020). The results show approximately 13,000 significant voxels covering 88 (of 489) different brain regions across all seven resting state networks as well as the brainstem, cerebellum, and the basal ganglia (Figure 3c; Table S3-4). We observed some voxel activations slightly outside the strict boundaries of gray matter, despite gray matter masking, which might be artifactual. Applying a more conservative gray matter mask excluded these voxels and led to a slight increase in performance (Δ*r =* .01).

Removing single regions or networks did not substantially impact the effect size in the hold-out sample (Figure 3d). The strongest reduction was observed when lesioning the somatomotor network, but the difference was still negligibly small (Δ*r* = .03). Hence, information widely distributed across the brain appears to be necessary to predict vulnerability to stress and does not rely on single networks or theory-driven regions.

#### Combining theory- and data-driven insights

Based on predictive success for the vulnerability facet in the scenes task, we explored associations with theory-driven neural targets using the same facet and task. There was a small significant correlation between vulnerability and amygdala responses (uncorrected), in agreement with common theories, but only when images were centered, as in the corresponding machine learning model (Figure 4). In contrast, dACC reactivity was associated with *lower* vulnerability to stress, contrary to the theory-guided direction, and was not affected by centering. Similar results were seen for the dorsal attention, frontoparietal, and default mode network. Lastly, there was increased reactivity of the sadness pattern in participants with higher vulnerability to stress. This is post-hoc evidence that some region-, network-, or signature-based measures might still warrant focused investigation for specific trait facets.

**Figure 4.**
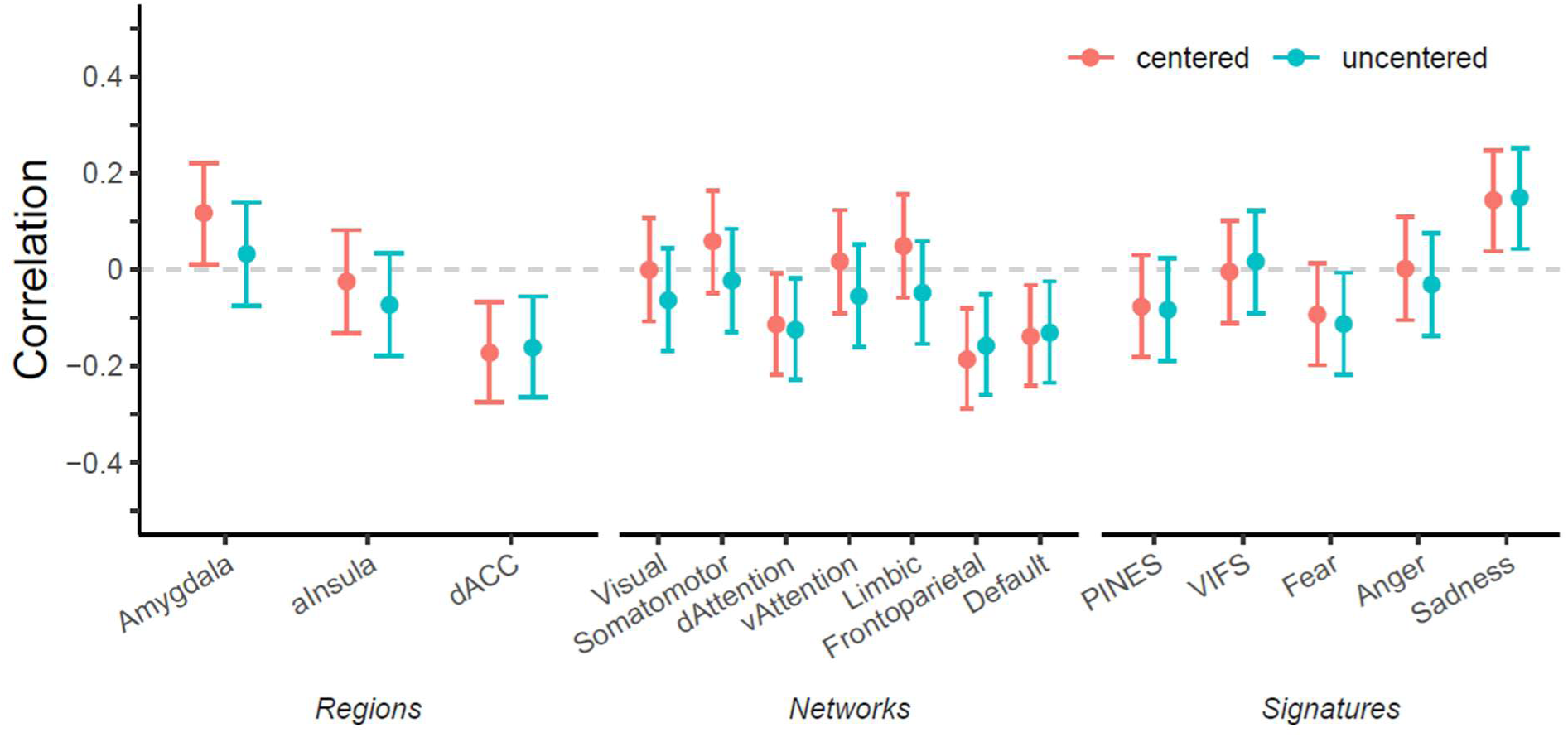
Correlations between theory-based neural targets and vulnerability to stress with versus without image-wise centering.

### Predictability of dynamic affective states

We tested how the predictability of individual differences in negative affective traits compares to the predictability of negative affective *states* based on dynamic self-reports collected during the scenes task. Using the same procedure as for the stress vulnerability pattern, the cross-validated within-person correlations averaged *r* = .88 (*SD* = .14; see supplemental methods). This means that when analyzing data from the same individual across different time points or conditions, the brain pattern exhibited a strong, consistent relationship with their momentary self-reported affect (Figure S1). Such high within-person correlations replicate previous findings on this dataset (r = .85; (Chang et al., 2015) and highlight the predictive gap between within-person and between-person prediction of affective constructs. Specifically, while neural patterns can robustly track fluctuations in an individual’s affective states over time, predicting stable individual differences (i.e., between-person effects) is substantially more challenging. As Figure S1 shows, this might be due to participants having considerably different offsets in their average brain activity regardless of stimulus intensity.

### Exploratory Multiverse Analysis

Figure 5 shows the cross-validated correlations of all models, separately for the different constructs serving as outcomes (see Figure 1c for all included modelling choices). Qualitatively, three important observations can be made. First, most neuroticism facets had predominantly positive cross-validated correlations, except for impulsiveness (Figure 5a). Secondly, vulnerability to stress had the strongest associations among neuroticism facets among a large range of models, in agreement with the results of the preregistered analyses. Thirdly, a comparison of different affective traits largely conforms to a face validity gradient (Figure 5b): The highest correlations were achieved for average task-based affective ratings, followed by negative affective dispositions (STAI, neuroticism other-reports, and neuroticism-self-reports), while positive affect had the lowest correlations, with depressive symptoms in between.

**Figure 5.**
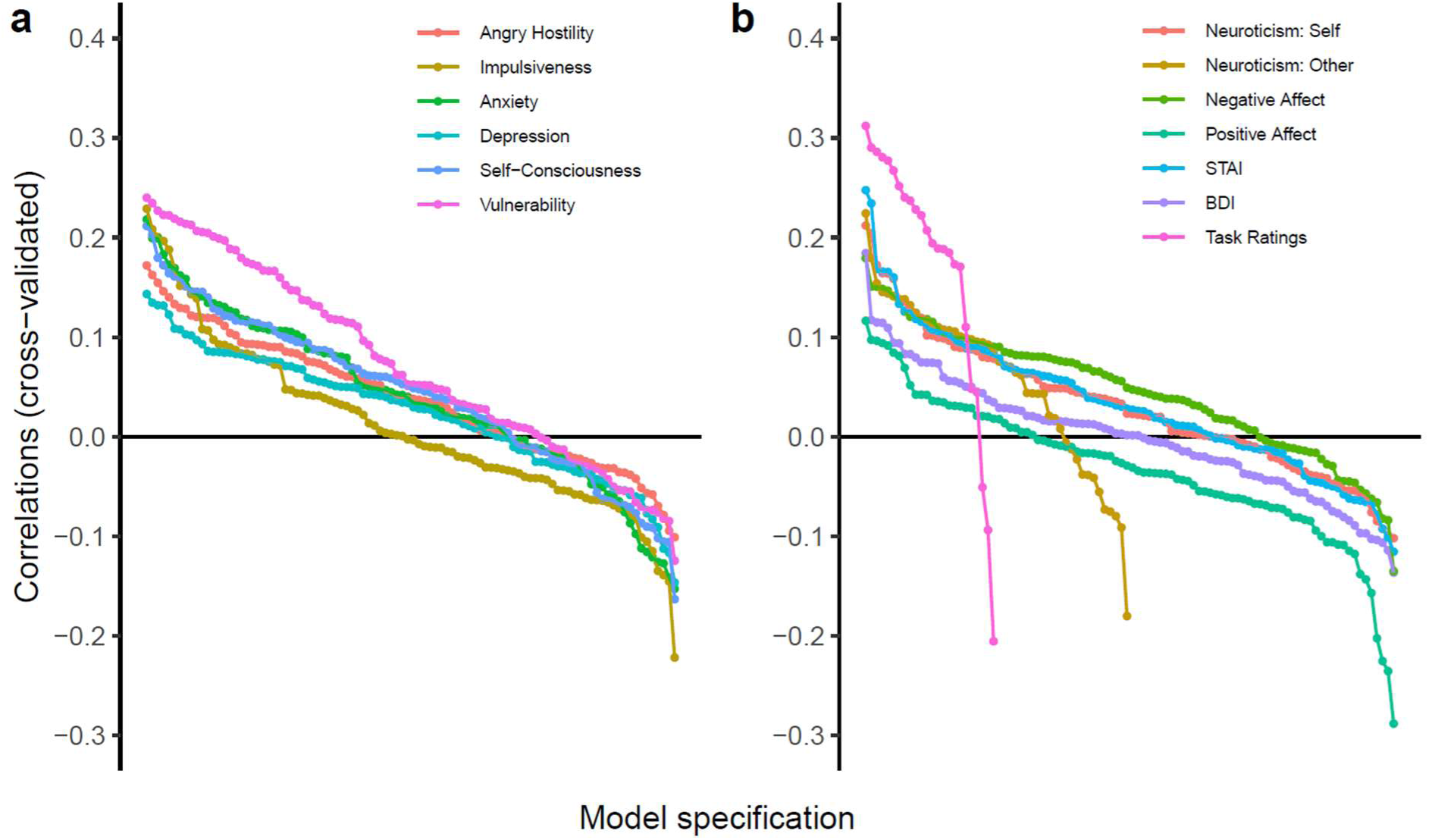
Cross-validated accuracy of all fitted models, separated by outcomes (a) neuroticism facets and (b) different affective traits. Models are ordered based on their accuracy along the x-axis. “Task ratings” refers to a participants’ average negative affective self-ratings during the scenes task.

A random forest regression on design factors explained 22% of variance in the size of correlations. The most important design factors were outcome choice and task choice, as confirmed by the variance decomposition (Figure 6). The scenes task provided overall better results than the faces task, potentially as it relates more closely to the experience rather than perception of emotion in others. This confirms that the two common fMRI tasks do not contain information on any trait, but that matching tasks and constructs is essential. Moreover, we found that PLS, PCR, and SVR worked similarly well, with worse performance for random forest regression. Also, using the full sample improved performance, supporting that higher accuracy might be achieved with larger samples. As there are a large number of design factor combinations which might be of interest for closer inspection, we developed an online application which provides an easy point-and-click user interface to interactively explore these results more closely: https://msicorello.shinyapps.io/ndesignshinyapp/.

**Figure 6.**
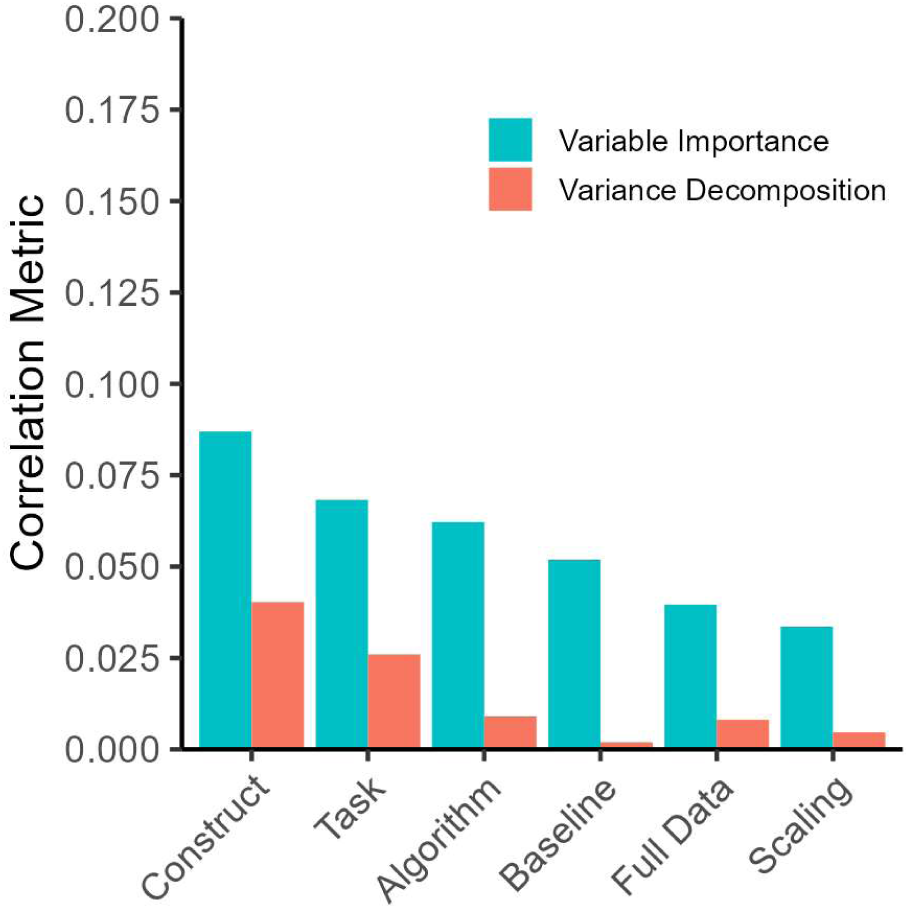
Design factors and cross-validated correlations. Blue bars show variable importance when predicting correlations from design factors using random forest regression. It reflects the drop in the accuracy when a given predictor is randomly shuffled in the metric of root mean squared error (i.e., average deviation between predicted and actual correlations). Red bars show standard deviations from a linear mixel model variance decomposition (i.e., how much correlations differ between design choices within a given factor).

## Discussion

Negative affectivity is a key dimension of well-being, personality, and psychopathology. Despite a large scientific literature, its neural basis is still unclear. Here, we presented the thus far most comprehensive investigation of the functional neurobiology underlying negative affectivity, combining the strengths of theory- and data-driven approaches across a large range of traits in the two most widely used affective tasks and supported by both preregistered and detailed exploratory analyses.

We found that the trait “vulnerability to stress” could be meaningfully predicted in the training sample. This association has high face validity and underlines the capability of task-based fMRI to predict specific negative affectivity traits. The predictive pattern was replicated in a hold-out sample and showed expected evidence of convergent and discriminant validity with other trait constructs, supporting both stability, reliability, and interpretability. Neurosynth-decoding and the absence of associations with specific affective neural signatures further supported these effects might be due to a domain-general neural responsiveness to stimulation in people with higher vulnerability to stress (e.g., instead of emotion-specific brain patterns). Hence, in contrast to broad neuroticism, a specific lower-order psychological trait like “stress vulnerability” appears to best match a relatively broad neuro-behavioral response of general sensory stimulation. This demonstrates the importance of identifying the correct alignment between the hierarchical granularity of psychological trait measures and neuro-behavioral measures (Brandt & Mueller, 2022). The resulting pattern was stable when common affect-related regions and networks were removed, showing that stress vulnerability is best represented by complex whole-brain patterns. Interestingly, the biggest performance drop was observed when removing the somato-motor network, which might support the perspective of emotion as an embodied experience (Reddan et al., 2024).

Similarly, the multiverse analysis suggests that negative affective traits differ in how well they can be predicted depending on their face validity for the fMRI task. For example, task-based self-ratings had the highest effects, followed by vulnerability to stress and trait anxiety, with the lowest effects for depressive symptoms and positive affect. Hence, great care should be taken in choosing the right target construct, being clear about the difference between nominal construct labels and actual construct meanings (Brandt & Mueller, 2022; Yarkoni, 2020).

For neuroticism as a broad trait domain of central importance, we found evidence *against* substantial theory-based effects of single brain regions, canonical resting state networks, or validated whole-brain signatures of negative affective states based on confidence intervals and Bayes factors. Similarly, a whole-brain machine learning approach did not lead to any predictive accuracy. The region-based analyses covered the amygdala, anterior insula, and dACC, which are commonly used in studies on well–being and psychopathology to attest affective problems. Hence, our results speak against the theory of a general amygdala hyper-responsiveness underlying neuroticism (Eysenck, 1967; Mitchell & Kumari, 2016) and simple psychological interpretations of activity in these regions. These results were stable even when using pattern-based predictive models for neural activity within these regions, which accounts for the fact that not all voxels within a region might have similar functional roles. Moreover, even data-driven regions most strongly correlated to neuroticism did not have significant associations in a split-half validation sample, with the best region identified from the scenes task even changing signs. Hence, a region-focused approach appears to be overall severely limited for the understanding of broad negative affectivity traits.

Similar observations were made for the network-based approaches, where the salience network usually received most attention, but had no substantial associations within our analyses, supported by Bayes factors. While the limbic and somatomotor networks did have nominally significant effects in the expected directions in the faces task, these effects did not survive the correction for multiple comparisons. Notably, we report Bayes factors without corrections for multiple comparisons, as different correction regiments might be sensible, and corrected Bayes factors would favor the null hypothesis even more strongly (Jong, 2019). Even accounting for the proposition that emotions require linear or non-linearly interacting combinations of network activities did not reveal statistically significant effects in multivariate predictive models (Lindquist & Barrett, 2012).

While these regions are still major targets in fMRI research on individual differences in emotional processes, null findings might be expected based on a process-oriented literature which demonstrated they are not specific to negative mental states, only weakly track task-based affective self-reports, and have very low test-retest reliabilities. In contrast, affective neural signatures might alleviate these problems, as they have much higher validity for affective states and acceptable test-retest reliabilities (Han et al., 2022; P. A. Kragel et al., 2018). Still, we found evidence against substantial associations between these neural signatures and neuroticism. As these neural signatures are well-validated for task-based affective self-ratings, this could indicate that the most common affective tasks do not probe affective traits on a self-report basis, even in the absence of neural measures. This is also reflected in the extremely small correlations between person-wise affective task ratings and negative affective traits (Table S1), which is a general problem in the field (Enkavi & Poldrack, 2020). This is further supported by our previous mega-analysis, which did not find hyperreactivity of affective neural signatures in participants with clinically elevated affective instability (Sicorello et al., 2021).

Importantly, when repeating the theory-driven models on the best data-driven model— predicting vulnerability to stress from image-wise centered brain data in the scenes task—a slightly different pattern emerged. Here, the amygdala showed the expected positive correlation with vulnerability, but only when images were centered on the person-mean. The opposite (theoretically unexpected) effect was observed for the dACC and three predominantly dorsal brain networks. In contrast to the amygdala, networks were not affected by centering. Hence, it is possible that global brain effects might bias amygdala reactivity towards negligibly small correlations, while removing this bias still leads to only small theory-congruent correlations due to its low reliability. Such a “whole-brain bias” is also supported by the offsets for within-person decoding of state-based self-reports and the observation that participants with higher activity in one resting state network, compared to other participants, were also much more likely to have higher activity in any other resting state network. This points towards a larger problem in fMRI for individual difference research and means successfully relating the amygdala to affective traits likely needs the right trait, the right task, the right preprocessing and at least several-hundred participants. Similarly, the correlation of vulnerability to stress with the neural signature for sadness, but not fear, anger or general negative affect, might indicate that it is crucial to test more closely which emotions are specifically elicited by common experimental emotion induction techniques.

For scientific progress on the neurobiology of negative affectivity, we believe that taking neuropsychometric principles seriously is essential. This necessitates improvements on both the psychological and biological side. On the psychological side, the most widely used affective task used in fMRI have been constructed to study affective *processes*, not individual differences. This results in tasks with low test-retest reliability and low construct validity even before neurobiology enters the picture, exemplified by the negligibly small correlations between average task ratings and negative affective traits or trait measures from other domains (Enkavi & Poldrack, 2020; Hedge et al., 2018; Schuch et al., 2022).

On the neurobiological side, there is also a general discrepancy in the predictive success of within- and between-person fMRI (Han et al., 2022; Jabakhanji et al., 2022). Even for our sample, affective task-based self-rating could be predicted with extremely high accuracy on a within-person state level (r = .88), reproducing previous findings (Chang et al., 2015). Despite this, accuracies for individual differences in task-based self-rating barely exceeded r = .30, which, notably, is still at the 75% percentile in non-biological individual difference research ((Gignac & Szodorai, 2016). Similar results have been found for physical pain (Han et al., 2022). Every trait-like questionnaire-based psychological construct is likely to have smaller effects than this ceiling, as they are affected by test-retest unreliability and have less face validity than task-based ratings. This was reflected in our multiverse analysis, which showed an effect size gradient along construct validity: Individual differences in task-based self ratings have the highest face validity and had the strongest effect sizes, followed by negative affective traits and trailed by positive affect, with depressivity in between.

Overall, some suggestions would be to (a) design tasks based on psychometric principles which show good test-retest reliability and valid associations between task behavior and questionnaires or daily life behavior, (b) leverage the properties of multivariate whole-brain patterns, especially in conjunction with hyperalignment (Haxby et al., 2020), (c) identify confounding sources of individual differences in whole-brain responsiveness or use image-wise scaling, (d) plan sample sizes for a noise ceiling between r = .20-.30 based on current fMRI approaches, expecting considerably smaller correlations, (e) leverage structural equation modeling combined with test-retest designs to increase effect sizes, and (f) aim to include the whole spectrum of trait realizations to avoid variance attenuation effects.

There are two notable limitations of this study. First, while the findings for the vulnerability pattern did replicate well in the hold-out sample, it is not clear whether results will generalize to other studies with different designs and sample compositions. Especially for clinical samples it should be noted that phenomena like trait dissociation can be important confounds, interfering with neuro-affective reactivity (Seitz et al., 2024). While our standard deviation in neuroticism is even slightly larger than in the representative validation sample by Costa and McCrae (1992), such studies often do not account for the representation of people with mental disorders. Accurately sampling (or even oversampling) such groups might lead to larger variation in negative affectivity and therefore larger effect sizes. Second, there might be physiological confounds which improved the predictive utility of our pattern. On the one hand, such confounds are not easily controlled, as there is a complex bi-directional relation between neural and physiological data (e.g., heart rate increases due to neural processing, but can also influence the fMRI signal). On the other hand, it might still be of interest to test the incremental value of our pattern above peripheral physiological measures in future studies.

In sum, our findings challenge conventional region- and network-based models of negative affectivity, demonstrating that individual differences therein are best captured by using distributed whole-brain patterns to predict more specific and fine-grained affective traits like stress vulnerability, whereas commonly targeted brain regions, networks, and affective signatures fail to provide reliable predictive power. These results highlight the necessity of aligning psychological constructs with neuro-behavioral measures at the appropriate level of granularity. By ensuring this match, future research can develop more stable and interpretable neural markers, advancing our understanding of affective well-being, personality, and dimensional models of mental health.

## References

Bates, S., Hastie, T., & Tibshirani, R. (2024). Cross-Validation: What Does It Estimate and How Well Does It Do It? Journal of the American Statistical Association, 119(546), 1434–1445. 10.1080/01621459.2023.2197686

Beck, A. T., Steer, R. A., & Brown, G. (1996). Beck Depression Inventory–II [Dataset]. 10.1037/t00742-000

Bolger, N., & Schilling, E. A. (1991). Personality and the Problems of Everyday Life: The Role of Neuroticism In Exposure and Reactivity to Daily Stressors. Journal of Personality, 59(3), 355–386. 10.1111/j.1467-6494.1991.tb00253.x

Bolger, N., & Zuckerman, A. (1995). Personality Processes and Individual Differences: A Framework for Studying Personality in the Stress Process. Journal of Personality and Social Psychology, 69(5), 890–902.

Brandt, A., & Mueller, E. M. (2022). Negative affect related traits and the chasm between self-report and neuroscience. Current Opinion in Behavioral Sciences, 43, 216–223. 10.1016/j.cobeha.2021.11.002

Casey, B. J., Cannonier, T., Conley, M. I., Cohen, A. O., Barch, D. M., Heitzeg, M. M., Soules, M. E., Teslovich, T., Dellarco, D. V., Garavan, H., Orr, C. A., Wager, T. D., Banich, M. T., Speer, N. K., Sutherland, M. T., Riedel, M. C., Dick, A. S., Bjork, J. M., Thomas, K. M.,… Dale, A. M. (2018). The Adolescent Brain Cognitive Development (ABCD) study: Imaging acquisition across 21 sites. Developmental Cognitive Neuroscience, 32, 43–54. 10.1016/j.dcn.2018.03.001

Čeko, M., Kragel, P. A., Woo, C.-W., López-Solà, M., & Wager, T. D. (2022). Common and stimulus-type-specific brain representations of negative affect. Nature Neuroscience, 25(6), 760–770. 10.1038/s41593-022-01082-w

Chang, L. J., Gianaros, P. J., Manuck, S. B., Krishnan, A., & Wager, T. D. (2015). A sensitive and specific neural signature for picture-induced negative affect. PLoS Biology, 13(6), 1–28. 10.1371/journal.pbio.1002180

Chen, G., Taylor, P. A., & Cox, R. W. (2017). Is the statistic value all we should care about in neuroimaging? NeuroImage, 147, 952–959. 10.1016/j.neuroimage.2016.09.066

Chen, Y.-W., & Canli, T. (2022). “Nothing to see here”: No structural brain differences as a function of the Big Five personality traits from a systematic review and meta-analysis. Personality Neuroscience, 5, e8. 10.1017/pen.2021.5

Costa Jr., P. T., & McCrae, R. R. (2008). The Revised NEO Personality Inventory (NEO-PI-R). In The SAGE handbook of personality theory and assessment, Vol 2: Personality measurement and testing (pp. 179–198). Sage Publications, Inc. 10.4135/9781849200479.n9

Costa, P. T., & McCrae, R. R. (1992). Revised NEO Personality Inventory (NEO-PI-R) and NEO Five-Factor Inventory (NEO-FFI) Professional Manual. Psychological Assessment Resources.

Crawford, J. R., & Henry, J. D. (2004). The Positive and Negative Affect Schedule (PANAS): Construct validity, measurement properties and normative data in a large non-clinical sample. British Journal of Clinical Psychology, 43(3), 245–265. 10.1348/0144665031752934

Cuijpers, P., Smit, F., Penninx, B. W. J. H., de Graaf, R., ten Have, M., & Beekman, A. T. F. (2010). Economic Costs of Neuroticism. Archives of General Psychiatry, 67(10), 1086. 10.1001/archgenpsychiatry.2010.130

Cunningham, W. A., & Brosch, T. (2012). Motivational salience: Amygdala tuning from traits, needs, values, and goals. Current Directions in Psychological Science, 21(1), 54–59. 10.1177/0963721411430832

Cuthbert, B. N. (2014). The RDoC framework: Facilitating transition from ICD/DSM to dimensional approaches that integrate neuroscience and psychopathology. World Psychiatry, 13(1), 28–35. 10.1002/wps.20087

DeYoung, C. G., Beaty, R. E., Genç, E., Latzman, R. D., Passamonti, L., Servaas, M. N., Shackman, A. J., Smillie, L. D., Spreng, R. N., Viding, E., & Wacker, J. (2022). Personality Neuroscience: An Emerging Field with Bright Prospects. Personality Science, 3, e7269. 10.5964/ps.7269

Diedrichsen, J., & Shadmehr, R. (2005). Detecting and adjusting for artifacts in fMRI time series data. NeuroImage, 27(3), 624–634. 10.1016/j.neuroimage.2005.04.039

Ekman, P., & Friesen, W. V. (1976). Pictures of Facial Affect. Consulting Psychologists Press.

Elliott, M. L., Knodt, A. R., Ireland, D., Morris, M. L., Poulton, R., Ramrakha, S., Sison, M. L., Moffitt, T. E., Caspi, A., & Hariri, A. R. (2020). What Is the Test-Retest Reliability of Common Task-Functional MRI Measures? New Empirical Evidence and a Meta-Analysis. Psychological Science, 31(7), 792–806. 10.1177/0956797620916786

Enkavi, A. Z., & Poldrack, R. A. (2020). Implications of the lacking relationship between cognitive task and self report measures for psychiatry. Biological Psychiatry: Cognitive Neuroscience and Neuroimaging. 10.1016/j.bpsc.2020.06.010

Eysenck, H. J. (1967). The biological basis of personality. Spring-field, Ill.

Fullana, M. A., Albajes-Eizagirre, A., Soriano-Mas, C., Vervliet, B., Cardoner, N., Benet, O., Radua, J., & Harrison, B. J. (2018). Fear extinction in the human brain: A meta-analysis of fMRI studies in healthy participants. Neuroscience and Biobehavioral Reviews, 88, 16–25. 10.1016/j.neubiorev.2018.03.002

Geuter, S., Reynolds Losin, E. A., Roy, M., Atlas, L. Y., Schmidt, L., Krishnan, A., Koban, L., Wager, T. D., & Lindquist, M. A. (2020). Multiple Brain Networks Mediating Stimulus–Pain Relationships in Humans. Cerebral Cortex, 30(7), 4204–4219. 10.1093/cercor/bhaa048

Gianaros, P. J., Kraynak, T. E., Kuan, D. C.-H., Gross, J. J., McRae, K., Hariri, A. R., Manuck, S. B., Rasero, J., & Verstynen, T. D. (2020). Affective brain patterns as multivariate neural correlates of cardiovascular disease risk. Social Cognitive and Affective Neuroscience, March, 1–12. 10.1093/scan/nsaa050

Gianaros, P. J., Marsland, A. L., Kuan, D. C.-H., Schirda, B. L., Jennings, J. R., Sheu, L. K., Hariri, A. R., Gross, J. J., & Manuck, S. B. (2014). An Inflammatory Pathway Links Atherosclerotic Cardiovascular Disease Risk to Neural Activity Evoked by the Cognitive Regulation of Emotion. Biological Psychiatry, 75(9), 738–745. 10.1016/j.biopsych.2013.10.012

Gignac, G. E., & Szodorai, E. T. (2016). Effect size guidelines for individual differences researchers. Personality and Individual Differences, 102, 74–78. 10.1016/j.paid.2016.06.069

Goodkind, M., Eickhoff, S. B., Oathes, D. J., Jiang, Y., Chang, A., Jones-Hagata, L. B., Ortega, B. N., Zaiko, Y. V., Roach, E. L., Korgaonkar, M. S., Grieve, S. M., Galatzer-Levy, I., Fox, P. T., & Etkin, A. (2015). Identification of a Common Neurobiological Substrate for Mental Illness. JAMA Psychiatry, 72(4), 305–315. 10.1001/jamapsychiatry.2014.2206

Gore, W. L., & Widiger, T. A. (2018). Negative emotionality across diagnostic models: RDoC, DSM-5 section III, and FFM. Personality Disorders: Theory, Research, and Treatment, 9(2), 155–164. 10.1037/per0000273

Han, X., Ashar, Y. K., Kragel, P., Petre, B., Schelkun, V., Atlas, L. Y., Chang, L. J., Jepma, M., Koban, L., Losin, E. A. R., Roy, M., Woo, C.-W., & Wager, T. D. (2022). Effect sizes and test-retest reliability of the fMRI-based neurologic pain signature. NeuroImage, 247, 118844. 10.1016/j.neuroimage.2021.118844

Haxby, J. V., Guntupalli, J. S., Nastase, S. A., & Feilong, M. (2020). Hyperalignment: Modeling shared information encoded in idiosyncratic cortical topographies. eLife, 9, 1–26. 10.7554/eLife.56601

Hedge, C., Powell, G., & Sumner, P. (2018). The reliability paradox: Why robust cognitive tasks do not produce reliable individual differences. Behavior Research Methods, 50(3), 1166–1186. 10.3758/s13428-017-0935-1

Inman, C. S., Bijanki, K. R., Bass, D. I., Gross, R. E., Hamann, S., & Willie, J. T. (2020). Human amygdala stimulation effects on emotion physiology and emotional experience. Neuropsychologia, February, 106722. 10.1016/j.neuropsychologia.2018.03.019

Jabakhanji, R., Vigotsky, A. D., Bielefeld, J., Huang, L., Baliki, M. N., Iannetti, G., & Apkarian, A. V. (2022). Limits of decoding mental states with fMRI. Cortex, 149, 101–122. 10.1016/j.cortex.2021.12.015

Jong, T. de. (2019). A Bayesian Approach to the Correction for Multiplicity. OSF. 10.31234/osf.io/s56mk

Kotov, R., Waszczuk, M. A., Krueger, R. F., Forbes, M. K., Watson, D., Clark, L. A., Achenbach, T. M., Althoff, R. R., Ivanova, M. Y., Michael Bagby, R., Brown, T. A., Carpenter, W. T., Caspi, A., Moffitt, T. E., Eaton, N. R., Forbush, K. T., Goldberg, D., Hasin, D., Hyman, S. E.,… Zimmerman, M. (2017). The hierarchical taxonomy of psychopathology (HiTOP): A dimensional alternative to traditional nosologies. Journal of Abnormal Psychology, 126(4), 454–477. 10.1037/abn0000258

Kozak, M. J., & Cuthbert, B. N. (2016). The NIMH Research Domain Criteria Initiative: Background, Issues, and Pragmatics. Psychophysiology, 53(3), 286–297. 10.1111/psyp.12518

Kragel, P. A., Koban, L., Barrett, L. F., & Wager, T. D. (2018). Representation, Pattern Information, and Brain Signatures: From Neurons to Neuroimaging. Neuron, 99(2), 257–273. 10.1016/j.neuron.2018.06.009

Kragel, P. A., & LaBar, K. S. (2015). Multivariate neural biomarkers of emotional states are categorically distinct. Social Cognitive and Affective Neuroscience, 10(11), 1437– 1448. 10.1093/scan/nsv032

Kragel, P., Han, X., Kraynak, T., Gianaros, P., & Wager, T. (2020). fMRI can be highly reliable, but it depends on what you measure (v2). PsyArXiv. 10.31234/osf.io/9eaxk

Krishnan, A., Woo, C.-W., Chang, L. J., Ruzic, L., Gu, X., López-Solà, M., Jackson, P. L., Pujol, J., Fan, J., & Wager, T. D. (2016). Somatic and vicarious pain are represented by dissociable multivariate brain patterns. eLife, 5, e15166. 10.7554/eLife.15166

Lang, P. J., Bradley, M. M., & Cuthbert, B. N. (2008). International affective picture system (IAPS): Affective ratings of pictures and instruction manual. Technical Report A-8. University of Florida, Gainesville, FL.

Lin, J., Li, L., Pan, N., Liu, X., Zhang, X., Suo, X., Kemp, G. J., Wang, S., & Gong, Q. (2023). Neural correlates of neuroticism: A coordinate-based meta-analysis of resting-state functional brain imaging studies. Neuroscience & Biobehavioral Reviews, 146, 105055. 10.1016/j.neubiorev.2023.105055

Lindquist, K. A., & Barrett, L. F. (2012). A functional architecture of the human brain: Emerging insights from the science of emotion. Trends in Cognitive Sciences, 16(11), 533–540. 10.1016/j.tics.2012.09.005

Lindquist, K. A., Satpute, A. B., Wager, T. D., Weber, J., & Barrett, L. F. (2016). The Brain Basis of Positive and Negative Affect: Evidence from a Meta-Analysis of the Human Neuroimaging Literature. Cerebral Cortex, 26(5), 1910–1922. 10.1093/cercor/bhv001

Liu, X., Lai, H., Li, J., Becker, B., Zhao, Y., Cheng, B., & Wang, S. (2021). Gray matter structures associated with neuroticism: A meta-analysis of whole-brain voxel-based morphometry studies. Human Brain Mapping, 42(9), 2706–2721. 10.1002/hbm.25395

Marek, S., Tervo-Clemmens, B., Calabro, F. J., Montez, D. F., Kay, B. P., Hatoum, A. S., Donohue, M. R., Foran, W., Miller, R. L., Hendrickson, T. J., Malone, S. M., Kandala, S., Feczko, E., Miranda-Dominguez, O., Graham, A. M., Earl, E. A., Perrone, A. J., Cordova, M., Doyle, O.,… Dosenbach, N. U. F. (2022). Reproducible brain-wide association studies require thousands of individuals. Nature, 603(7902), 654–660. 10.1038/s41586-022-04492-9

McCrae, R. R., & Costa Jr., P. T. (2008). The five-factor theory of personality. In Handbook of personality: Theory and research, 3rd ed (pp. 159–181). The Guilford Press.

Miller, K. L., Alfaro-Almagro, F., Bangerter, N. K., Thomas, D. L., Yacoub, E., Xu, J., Bartsch, A. J., Jbabdi, S., Sotiropoulos, S. N., Andersson, J. L., Griffanti, L., Douaud, G., Okell, T. W., Weale, P., Dragonu, I., Garratt, S., Hudson, S., Collins, R., Jenkinson, M.,… Smith, S. M. (2016). Multimodal population brain imaging in the UK Biobank prospective epidemiological study. Nature Neuroscience, 19(11), 1523– 1536. 10.1038/nn.4393

Mincic, A. M. (2015). Neuroanatomical correlates of negative emotionality-related traits: A systematic review and meta-analysis. Neuropsychologia, 77, 97–118. 10.1016/j.neuropsychologia.2015.08.007

Mitchell, R. L. C., & Kumari, V. (2016). Hans Eysenck’s interface between the brain and personality: Modern evidence on the cognitive neuroscience of personality. Personality and Individual Differences, 103, 74–81. 10.1016/j.paid.2016.04.009

Morey, R. D., & Rouder, J. N. (2018). BayesFactor: Computation of Bayes Factors for Common Designs. R package version 0.9.12-4.2 [Computer software]. https://cran.r-project.org/package=BayesFactor

Noble, S., Scheinost, D., & Constable, R. T. (2019). A decade of test-retest reliability of functional connectivity: A systematic review and meta-analysis. NeuroImage, 203(August), 116157. 10.1016/j.neuroimage.2019.116157

Ormel, J., Bastiaansen, A., Riese, H., Bos, E. H., Servaas, M., Ellenbogen, M., Rosmalen, J. G. M., & Aleman, A. (2013). The biological and psychological basis of neuroticism: Current status and future directions. Neuroscience & Biobehavioral Reviews, 37(1), 59–72. 10.1016/j.neubiorev.2012.09.004

Ormel, J., Jeronimus, B. F., Kotov, R., Riese, H., Bos, E. H., Hankin, B., Rosmalen, J. G. M., & Oldehinkel, A. J. (2013). Neuroticism and common mental disorders: Meaning and utility of a complex relationship. Clinical Psychology Review, 33(5), 686–697. 10.1016/j.cpr.2013.04.003

Reddan, M. C., Chang, L., Kragel, P., & Wager, T. D. (2024). Somatosensory and motor contributions to emotion representation (arXiv:2411.08973). arXiv. 10.48550/arXiv.2411.08973

Ringwald, W. R., Nielsen, S. R., Mostajabi, J., Vize, C. E., van den Berg, T., Manuck, S. B., Marsland, A. L., & Wright, A. G. C. (2024). Characterizing stress processes by linking big five personality states, traits, and day-to-day stressors. Journal of Research in Personality, 110, 104487. 10.1016/j.jrp.2024.104487

Schuch, S., Philipp, A. M., Maulitz, L., & Koch, I. (2022). On the reliability of behavioral measures of cognitive control: Retest reliability of task-inhibition effect, task-preparation effect, Stroop-like interference, and conflict adaptation effect. Psychological Research, 86(7), 2158–2184. 10.1007/s00426-021-01627-x

Schulz, M.-A., Bzdok, D., Haufe, S., Haynes, J.-D., & Ritter, K. (2024). Performance reserves in brain-imaging-based phenotype prediction. Cell Reports, 43(1). 10.1016/j.celrep.2023.113597

Schulze, L., Schulze, A., Renneberg, B., Schmahl, C., & Niedtfeld, I. (2019). Neural Correlates of Affective Disturbances: A Comparative Meta-analysis of Negative Affect Processing in Borderline Personality Disorder, Major Depressive Disorder, and Posttraumatic Stress Disorder. Biological Psychiatry: Cognitive Neuroscience and Neuroimaging, 4(3), 220–232. 10.1016/j.bpsc.2018.11.004

Seeley, W. W. (2019). The Salience Network: A Neural System for Perceiving and Responding to Homeostatic Demands. Journal of Neuroscience, 39(50), 9878–9882. 10.1523/JNEUROSCI.1138-17.2019

Seitz, K. I., Sicorello, M., Schmitz, M., Valencia, N., Herpertz, S. C., Bertsch, K., & Neukel, C. (2024). Childhood Maltreatment and Amygdala Response to Interpersonal Threat in a Transdiagnostic Adult Sample: The Role of Trait Dissociation. Biological Psychiatry: Cognitive Neuroscience and Neuroimaging, 9(6), 626–634. 10.1016/j.bpsc.2024.01.003

Servaas, M. N., van der Velde, J., Costafreda, S. G., Horton, P., Ormel, J., Riese, H., & Aleman, A. (2013). Neuroticism and the brain: A quantitative meta-analysis of neuroimaging studies investigating emotion processing. Neuroscience and Biobehavioral Reviews, 37(8), 1518–1529. 10.1016/j.neubiorev.2013.05.005

Sha, Z., Wager, T. D., Mechelli, A., & He, Y. (2019). Common Dysfunction of Large-Scale Neurocognitive Networks Across Psychiatric Disorders. Biological Psychiatry, 85(5), 379–388. 10.1016/j.biopsych.2018.11.011

Shackman, A. J., Tromp, D. P. M., Stockbridge, M. D., Kaplan, C. M., Tillman, R. M., & Fox, A. S. (2016). Dispositional negativity: An integrative psychological and neurobiological perspective. Psychological Bulletin, 142(12), 1275–1314. 10.1037/bul0000073

Sicorello, M., Herzog, J., Wager, T. D., Ende, G., Müller-Engelmann, M., Herpertz, S. C., Bohus, M., Schmahl, C., Paret, C., & Niedtfeld, I. (2021). Affective neural signatures do not distinguish women with emotion dysregulation from healthy controls: A mega-analysis across three task-based fMRI studies. Neuroimage: Reports, 1(2), 100019. 10.1016/j.ynirp.2021.100019

Sicorello, M., & Schmahl, C. (2021). Emotion dysregulation in borderline personality disorder: A fronto–limbic imbalance? Current Opinion in Psychology, 37, 114–120. 10.1016/j.copsyc.2020.12.002

Silverman, M. H., Wilson, S., Ramsay, I. S., Hunt, R. H., Thomas, K. M., Krueger, R. F., & Iacono, W. G. (2019). Trait Neuroticism and Emotion Neurocircuitry: fMRI Evidence for a Failure in Emotion Regulation. Development and Psychopathology, 31(3), 1085– 1099. 10.1017/S0954579419000610

Simonsohn, U., Simmons, J. P., & Nelson, L. D. (2020). Specification curve analysis. Nature Human Behaviour, 4(11), 1208–1214. 10.1038/s41562-020-0912-z

Spielberger, C. D. (1989). State-Trait Anxiety Inventory: Bibliography (2nd ed.). Consulting Psychologists Press.

Szucs, D., & Ioannidis, J. P. A. (2017). Empirical assessment of published effect sizes and power in the recent cognitive neuroscience and psychology literature. PLoS Biology, 15(3), 1–18. 10.1371/journal.pbio.2000797

Taschereau-Dumouchel, V., Kawato, M., & Lau, H. (2019). Multivoxel pattern analysis reveals dissociations between subjective fear and its physiological correlates. Molecular Psychiatry. 10.1038/s41380-019-0520-3

Thomas Yeo, B. T., Krienen, F. M., Sepulcre, J., Sabuncu, M. R., Lashkari, D., Hollinshead, M., Roffman, J. L., Smoller, J. W., Zöllei, L., Polimeni, J. R., Fisch, B., Liu, H., & Buckner, R. L. (2011). The organization of the human cerebral cortex estimated by intrinsic functional connectivity. Journal of Neurophysiology, 106(3), 1125–1165. 10.1152/jn.00338.2011

Uddin, L. Q. (2015). Salience processing and insular cortical function and dysfunction. Nature Reviews Neuroscience, 16(1), 55–61. 10.1038/nrn3857

Visser, R. M., Bathelt, J., Scholte, H. S., & Kindt, M. (2021). Robust BOLD Responses to Faces But Not to Conditioned Threat: Challenging the Amygdala’s Reputation in Human Fear and Extinction Learning. The Journal of Neuroscience, 41(50), 10278– 10292. 10.1523/JNEUROSCI.0857-21.2021

Wager, T. D. (2024). CanlabCore [Computer software]. https://github.com/canlab/CanlabCore

Widaman, K. F., & Revelle, W. (2023). Thinking thrice about sum scores, and then some more about measurement and analysis. Behavior Research Methods, 55(2), 788–806. 10.3758/s13428-022-01849-w

Wright, A. G. C., & Simms, L. J. (2015). A metastructural model of mental disorders and pathological personality traits. Psychological Medicine, 45(11), 2309–2319. 10.1017/S0033291715000252

Xu, J., Van Dam, N. T., Feng, C., Luo, Y., Ai, H., Gu, R., & Xu, P. (2019). Anxious brain networks: A coordinate-based activation likelihood estimation meta-analysis of resting-state functional connectivity studies in anxiety. Neuroscience & Biobehavioral Reviews, 96, 21–30. 10.1016/j.neubiorev.2018.11.005

Yarkoni, T. (2020). The generalizability crisis. Behavioral and Brain Sciences. 10.1017/S0140525X20001685

Yarkoni, T., Poldrack, R. A., Nichols, T. E., Van Essen, D. C., & Wager, T. D. (2011). Large-scale automated synthesis of human functional neuroimaging data. Nature Methods, 8(8), 665–670. 10.1038/nmeth.1635

Zhou, F., Zhao, W., Qi, Z., Geng, Y., Yao, S., Kendrick, K. M., Wager, T. D., & Becker, B. (2021). A distributed fMRI-based signature for the subjective experience of fear. Nature Communications, 12(1), 1–16. 10.1038/s41467-021-26977-3

Zugman, A., Jett, L., Antonacci, C., Winkler, A. M., & Pine, D. S. (2023). A systematic review and meta-analysis of resting-state fMRI in anxiety disorders: Need for data sharing to move the field forward. Journal of Anxiety Disorders, 99, 102773. 10.1016/j.janxdis.2023.102773

