## Supplemental Materials for "The functional neurobiology of negative affective traits across regions, networks, signatures, and a machine learning multiverse"

*-Supplements-*

### MRI data acquisition and preprocessing

Functional BOLD image acquisition parameters for the facial-expression tasks were: field-of-view (FOV) = 200×200mm, matrix size = 64×64, time-to-repetition (TR) = 2000ms, time-to-echo (TE) = 29ms, and flip angle (FA) = 90°. Thirty-four slices per volume were collected along an inferior-to-superior encoding direction, with each volume having a 3mm thickness and no gap. A total of 195 and 273 BOLD signal volumes were collected throughout the facial-expression tasks using PFA and NIM-STIM images, respectively. Functional BOLD image acquisition parameters for the IAPS task were: FOV = 205×205mm, matrix size = 64×64, TR = 2000ms, TE = 28ms, and FA = 90°. The entire task duration was 11 min and 16 sec (15 'Look neutral' trials; 15 'Look negative' trials; 15 'Decrease negative' trials). Images were presented such that no more than 2 identical trial types ("Look negative" or "Decrease negative") were consecutive, and no more than 4 unpleasant images were consecutive. Thirtynine slices per volume were collected along an inferior-to-superior encoding direction, with each volume having a 3mm thickness and no gap (338 task volumes in total). A 6-sec countdown preceded task onset. The 3 volumes of this countdown were not modeled in the design matrix, nor were remaining 3 volumes collected after the offset of the final rest period (344 functional run volumes in total). To co-register BOLD signal volumes, T1-weighted magnetization prepared rapid gradient echo (MPRAGE) images were acquired over 7 min and 17 sec by these parameters: FOV = 256×208mm, matrix size = 256×208, TR = 2100ms, inversion time (TI) = 1100ms, TE = 3.31ms, and FA = 8° (192 slices, 1mm thickness, no gap).

All fMRI data for the facial-expression and IAPS tasks were preprocessed with statistical parametric mapping software (SPM12; http://www.fil.ion.ucl.ac.uk/spm). For spatial preprocessing, T1-weighted MPRAGE images were classified into 6 tissue types. Biasedcorrected and deformation field maps were then computed. Functional images were realigned to the first image of the series by 6-parameter rigid-body transformation, using the re-slice step to match the first image on a voxel-by-voxel basis. Before realignment, slice-timing correction was applied to IAPS fMRI task data to account for acquisition time variation in this eventrelated design. Realigned images were co-registered to each participant's skull-stripped and biased-corrected MPRAGE image. Co-registered images were then normalized to Montreal Neurological Institute (MNI) space. Normalized images were then smoothed by a 6mm fullwidth-at-half-maximum (FWHM) Gaussian kernel.

### Details on Machine Larning Algorithms

Model performance in the training sample was estimated using 2x5x5 nested cross-validation the inner folds being used for hyperparameter optimization, and the outer folds for independent testing using the chosen optimal hyperparameter (PLS/PCR: number of components between one and the sample size of the innermost training fold optimized using Bayesian parameter optimization, SVM: Slack parameter C optimized using grid search, RF: mtry with grid search between one and total number of variables after dimensionality reduction using PCA with the number of components equal to the sample size of the innermost training fold). The procedure was repeated two times for more stability. Product-moment correlations were used as a performance metric, as the aim was to train a pattern for scale-independent correlates of negative affectivity, which can be used across different studies with different scanning parameters or different questionnaires. The resulting 10 product-moment correlations (two repeats times five outer folds) were Fisher-transformed, averaged, and the average back-transformed for the final model estimate. The model configuration with the highest average correlation was chosen for evaluation in the hold-out sample. Before evaluation, the model was retrained on the full training data using an optimal hyperparameter chosen based on 5-fold cross-validation (corresponding to the inner folds of the nested procedure). Using this final model, neuroticism scores of the hold-out sample were correlated with model predictions, generated by taking the dot product between machine learning pattern and fMRI data. This final correlation was supplemented with p-values and Bayes factors, both one-sided.

**Method for within-person prediction**

Within-person prediction had been previously performed on the same dataset for the creation of the PINES signature by Chang and colleagues (2015; see main manuscript). They prepared one beta image for each person and rating to a negative picture, i.e., as ratings varied between 1 and 5, each participant is associated with five beta images. These images are openly available on neurovault: https://neurovault.org/collections/1964/. In accordance with the previous study, we stacked the images on top of each other in a single data matrix. Then, we performed 10-fold cross-validation, leaving out 10% of participants in each iteration to avoid dependencies in training and test data. Nine folds were used for training a predictive model based on the specifications of the best model we identified for individual differences, i.e., support vector regression after image-wise centering. In the remaining test fold, a correlation between actual and predicted ratings was calculated separately for each participant. This procedure was repeated until each participant was part of a test fold, providing person-specific correlations for the whole sample. We report the mean value and standard deviation for these person-wise correlations in the main paper.

**Table S1**

*Descriptive statistics for psychological traits*

| Construct | *M* | *SD* | 1 | 2 | 3 | 4 | 5 | 6 | 7 | 8 | 9 | 10 | 11 | 12 | 13 |
|---|---|---|---|---|---|---|---|---|---|---|---|---|---|---|---|
| 1. Neuroticism | 75.5 | 21.7 | | | | | | | | | | | | | |
| 2. N1: Anxiety | 13.7 | 5.2 | 0.81 | | | | | | | | | | | | |
| 3. N2: Hostility | 12.0 | 5.0 | 0.74 | 0.52 | | | | | | | | | | | |
| 4. N3: Depression | 11.5 | 5.6 | 0.84 | 0.63 | 0.54 | | | | | | | | | | |
| 5. N4: Self-Consciousness | 13.8 | 4.5 | 0.71 | 0.55 | 0.36 | 0.55 | | | | | | | | | |
| 6. N5: Impulsiveness | 15.4 | 4.6 | 0.59 | 0.34 | 0.39 | 0.35 | 0.25 | | | | | | | | |
| 7. N6: Vulnerability | 9.1 | 4 | 0.80 | 0.6 | 0.51 | 0.67 | 0.52 | 0.37 | | | | | | | |
| 8. N: Other-Report | 16.2 | 9.1 | 0.42 | 0.36 | 0.37 | 0.36 | 0.27 | 0.19 | 0.34 | | | | | | |
| 9. Negative Affect | 15.8 | 5.3 | 0.62 | 0.53 | 0.45 | 0.62 | 0.38 | 0.30 | 0.48 | 0.29 | | | | | |
| 10. STAI | 32.5 | 8.4 | 0.71 | 0.58 | 0.49 | 0.71 | 0.47 | 0.30 | 0.63 | 0.39 | 0.59 | | | | |
| 11. BDI | 4.2 | 4.4 | 0.44 | 0.31 | 0.32 | 0.50 | 0.26 | 0.23 | 0.36 | 0.33 | 0.45 | 0.62 | | | |
| 12. Positive Affect | 34.5 | 6.5 | -0.33 | -0.21 | -0.22 | -0.32 | -0.28 | -0.12 | -0.37 | -0.13 | -0.11 | -0.37 | -0.29 | | |
| 13. Extraversion | 115 | 19.5 | -0.33 | -0.23 | -0.18 | -0.31 | -0.39 | -0.04 | -0.36 | -0.11 | -0.21 | -0.30 | -0.17 | 0.49 | |
| 14. Task-Ratings | 2.4 | 0.7 | 0.03 | 0.10 | -0.03 | -0.01 | 0.12 | -0.07 | 0.03 | -0.02 | 0.02 | -0.03 | -0.10 | -0.02 | 0.09 |

*Note.* STAI = State-Trait Anxiety Inventory (trait scores). BDI = Beck Depression Inventory. Task-Ratings = Person-wise average difference between affective task ratings between negative and neutral scenes. Sample sizes of correlations can differ due to missingness and study-related factors (e.g., many participants only performed the faces task, which does not include task-based ratings, and many participants did not contribute other reports for neuroticism)

**Table S2**

*Theory-driven neural associations with neuroticism*

| | IAPS ($N$ = 332) | | | FACES ($N$ = 424) | | |
|---|---|---|---|---|---|---|
| | *r* | 95% CI | $BF_{01}$ | r | 95% CI | $BF_{01}$ |
| | | | *Regions* | | | |
| *Averaged signal* | | | | | | |
| Amygdala-L | .03 | [-.08, .14] | 19.4 | .07 | [-.03, .16] | 10.5 |
| Amygdala-R | .05 | [-.06, .15] | 16.4 | .06 | [-.04, .15] | 12.6 |
| aInsula-L | -.05 | [-.15, .06] | 16.1 | .03 | [-.06, .13] | 20.4 |
| aInsula-R | -.06 | [-.17, .05] | 12.7 | .05 | [-.05, .14] | 16.6 |
| dACC-L | -.11 | [-.22, .00] | 3.4 | .04 | [-.06, .13] | 18.7 |
| dACC-R | -.11 | [-.22, -.00] | 2.84 | .04 | [-.05, .13] | 17.0 |
| *PLS Models*[a] | | | | | | |
| Amygdala | -.05 | [-.24, .15] | 11.5 | .13 | [-.06, .32] | 5.3 |
| aInsula | -.05 | [-.24, .15] | 11.4 | .02 | [-.18, .21] | 12.6 |
| dACC | .04 | [-.16, .23] | 11.9 | .09 | [-.11, .28] | 8.6 |
| *Best region* | | | | | | |
| Cerebellum/ Diencephalon | -.12 | [-.26, .03] | 4.95 | .07 | [-.06, .21] | 9.5 |
| | | | *Networks* | | | |
| *Averaged Signal* | | | | | | |
| Visual | -.02 | [-.13, .09] | 21.6 | .04 | [-.06, .13] | 19.4 |
| Somatomotor | -.02 | [-.12, .09] | 21.9 | .11 | [.02, .20] | 1.9 |
| dAttention | -.03 | [-.13, .08] | 20.5 | .07 | [-.03, .16] | 9.5 |
| vAttention | -.04 | [-.15, .07] | 17.5 | .07 | [-.03, .16] | 9.9 |
| Limbic | -.01 | [-.12, .09] | 22.2 | .12 | [.03, .22] | 1.0 |
| Frontoparietal | -.10 | [-.20, .01] | 4.7 | .04 | [-.05, .14] | 17.0 |
| Default-Mode | -.07 | [-.18, .04] | 10.1 | .06 | [-.04, .15] | 12.5 |
| *Linear Regression* | | | | | | |
| All networks[b] | -.03 | .467 | >100 | .12 | .071 | >100 |
| *Random Forest* | | | | | | |
| All networks[b] | .10 | .059 | | -.06 | .710 | |
| | | | *Signatures* | | | |
| PINES | -.09 | [-.20, 0.02] | 5.5 | .07 | [-.03, .16] | 9.5 |
| VIFS | .04 | [-.07, .15] | 17.6 | -.03 | [-.13, .07] | 21.3 |
| Fear | .05 | [-.06, .15] | 16.2 | .01 | [-.09, .10] | 25.5 |
| Anger | .01 | [-.10, .12] | 22.4 | .05 | [-.05, .14] | 16.1 |
| Sadness | -.03 | [-.13, .08] | 20.6 | .05 | [-.05, .14] | 15.4 |

Note. $BF_{01}$ = Bayes factor of the null hypothesis over the alternative hypothesis. Bayes factors were calculated with the bayesFactor toolbox in matlab.

[a]The proportion of in-region voxels weights with positive signs ranged between 49-51%.

[b]For the linear multiple regression approach, the multiple correlation coefficient is reported, calculated as the square-root of adjusted $R^2$. Here, $p$-values are reported instead of confidence intervals, as confidence intervals for (unadjusted) $R^2$ cannot contain zero and are therefore harder to interpret in terms of statistical significance. Similarly, $p$-values are reported for the random forest approach, as it was preregistered to be conducted on the whole sample with out-of-bag prediction for which (to our knowledge) currently are no valid confidence intervals available. The $p$-value is for a one-sided test, as negative correlations between predictions and actual values are not meaningful.

**Table S3**

*Clusters of voxels with positive weights at FDR of q = .10*

| Region | Volume [mm³] | MNI | | | $Z_{max}$-Statistic | % covered |
|---|---|---|---|---|---|---|
| | | X | Y | Z | | |
| Basal ganglia | 64 | -12 | -14 | -4 | 3.652305768 | 100 |
| Basal ganglia | 64 | -12 | 10 | 12 | 3.852284309 | 100 |
| Basal ganglia | 144 | 32 | -32 | 0 | 3.90804411 | 89 |
| Basal ganglia | 160 | -24 | -16 | 8 | 4.104912286 | 100 |
| Brainstem | 224 | 10 | -24 | -34 | 4.04452717 | 79 |
| Brainstem | 224 | -2 | -26 | -12 | 3.983246299 | 43 |
| Brainstem | 472 | -4 | -10 | -22 | 4.454181836 | 32 |
| Cerebellum | 64 | 28 | -56 | -42 | 3.545403913 | 25 |
| Cerebellum | 64 | 8 | -44 | -42 | 3.73168988 | 100 |
| Cerebellum | 64 | 38 | -74 | -38 | 3.666787978 | 88 |
| Cerebellum | 64 | -22 | -60 | -36 | 3.601441673 | 88 |
| Cerebellum | 128 | 22 | -54 | -36 | 4.052773897 | 75 |
| Cerebellum | 144 | 34 | -68 | -40 | 3.96928996 | 67 |
| Cerebellum | 144 | 12 | -48 | -16 | 4.026818128 | 56 |
| Cerebellum | 208 | 18 | -70 | -14 | 4.175139382 | 62 |
| Cerebellum | 416 | 12 | -66 | -38 | 3.895829671 | 54 |
| Cerebellum | 776 | -12 | -68 | -38 | 4.334853742 | 35 |
| Cortex Default ModeA | 128 | -22 | 20 | 44 | 3.605726765 | 50 |
| Cortex Default ModeA | 144 | -8 | 62 | 8 | 3.953326055 | 100 |
| Cortex Default ModeB | 64 | -58 | -46 | -2 | 3.681519856 | 50 |
| Cortex Default ModeB | 128 | -42 | 24 | -14 | 3.650804356 | 75 |
| Cortex Default ModeB | 208 | 16 | 48 | 26 | 4.272855498 | 35 |
| Cortex Default ModeC | 64 | 34 | -26 | -24 | 3.566796028 | 63 |
| Cortex Default ModeC | 200 | -38 | -26 | -18 | 3.786977859 | 52 |
| Cortex Dorsal AttentionA | 256 | 28 | -60 | 30 | 4.303589365 | 41 |
| Cortex Dorsal AttentionA | 392 | 24 | -78 | 20 | 3.996921822 | 33 |
| Cortex Dorsal AttentionA | 432 | 40 | -42 | 66 | 4.388786267 | 13 |
| Cortex Dorsal AttentionB | 64 | 12 | -54 | 60 | 3.927093939 | 100 |
| Cortex Dorsal AttentionB | 64 | -42 | -52 | 62 | 3.646879984 | 75 |
| Cortex Dorsal AttentionB | 216 | -38 | -4 | 32 | 4.343334203 | 7 |
| Cortex Dorsal AttentionB | 296 | -32 | -54 | 68 | 4.411285513 | 84 |
| Cortex Fronto ParietalA | 128 | 28 | 14 | 28 | 4.304903366 | 25 |
| Cortex Fronto ParietalB | 128 | -58 | -50 | -16 | 3.943971404 | 100 |
| Cortex Fronto ParietalB | 288 | 64 | -20 | -18 | 3.843402842 | 75 |
| Cortex Fronto ParietalC | 256 | 0 | -74 | 44 | 3.597457891 | 50 |
| Cortex Fronto ParietalC | 432 | -2 | -72 | 52 | 4.254097528 | 43 |
| Cortex Limbic | 64 | 18 | -4 | -42 | 3.529895563 | 25 |
| Cortex Limbic | 128 | 60 | -16 | -34 | 3.779750831 | 63 |
| Cortex Limbic | 160 | 28 | 0 | -38 | 3.806306957 | 55 |
| Cortex Limbic | 176 | 44 | 20 | -20 | 4.221152529 | 73 |
| Cortex Limbic | 176 | 16 | 52 | -18 | 3.893277031 | 68 |

| | | | | | | |
|---|---|---|---|---|---|---|
| Cortex Limbic | 288 | 54 | -12 | -36 | 4.256764093 | 89 |
| Cortex Limbic | 320 | 54 | 8 | -32 | 3.992710966 | 98 |
| Cortex Limbic | 424 | 22 | -8 | -44 | 4.367441661 | 28 |
| Cortex Limbic | 520 | -28 | 0 | -40 | 4.086828182 | 66 |
| Cortex Limbic | 736 | 36 | 20 | -38 | 5.03981465 | 97 |
| Cortex SomatomotorB | 600 | 40 | -18 | 2 | 3.833716071 | 28 |
| Cortex Temporal Parietal | 256 | -48 | 6 | -6 | 4.158587209 | 41 |
| Cortex Ventral AttentionA | 192 | 42 | -4 | -4 | 4.009962572 | 79 |
| Cortex Ventral AttentionA | 320 | 38 | -14 | -10 | 4.447322313 | 25 |
| Cortex Ventral AttentionB | 64 | -42 | 46 | 28 | 3.958137593 | 100 |
| Cortex Ventral AttentionB | 224 | 48 | 38 | 14 | 4.141441415 | 21 |
| Cortex Ventral AttentionB | 272 | 2 | 32 | 18 | 4.040992933 | 82 |
| Cortex Visual Central | 64 | 40 | -68 | -20 | 3.586643375 | 50 |
| Cortex Visual Central | 128 | -16 | -88 | 12 | 3.842890959 | 6 |
| Cortex Visual Central | 336 | 2 | -84 | 22 | 3.910583629 | 38 |
| Cortex Visual Peripheral | 64 | -8 | -70 | -2 | 3.536887002 | 100 |
| Cortex Visual Peripheral | 64 | -22 | -74 | 2 | 3.534790704 | 50 |
| Cortex Visual Peripheral | 64 | -24 | -86 | 36 | 3.593700695 | 75 |
| Cortex Visual Peripheral | 384 | 8 | -62 | 6 | 4.277608272 | 83 |
| Cortex Visual Peripheral | 552 | 26 | -64 | 6 | 4.035739112 | 49 |
| Diencephalon | 64 | 18 | -24 | -4 | 3.57039058 | 100 |
| No description | 64 | 68 | -48 | 4 | 3.524026159 | 0 |
| No description | 64 | 24 | 22 | 10 | 3.547524833 | 0 |
| No description | 64 | 28 | -58 | 12 | 3.588740176 | 0 |
| No description | 64 | 28 | -16 | 52 | 3.526708569 | 0 |
| No description | 64 | -22 | -76 | 58 | 4.050048247 | 0 |
| No description | 128 | 44 | -40 | 28 | 4.08176954 | 0 |

*Note.* The "% covered" column indicates the percent of voxels in a cluster cover by a specific region label. Automated labelling was performed with the canlab toolbox using the region() and table() functions.

**Table S4**

*Clusters of voxels with negative weights at FDR of q = .10*

| Region | Volume [mm³] | MNI | | | $Z_{max}$-Statistic | % covered |
|---|---|---|---|---|---|---|
| | | X | Y | Z | | |
| Basal ganglia | 64 | 6 | 0 | -10 | -3.54718 | 75 |
| Basal ganglia | 64 | 2 | 12 | 16 | -3.7611 | 50 |
| Basal ganglia | 88 | 0 | 14 | 14 | -3.55865 | 9 |
| Basal ganglia | 144 | -24 | 12 | -4 | -3.80977 | 72 |
| Basal ganglia | 264 | -8 | 0 | 18 | -4.20653 | 100 |
| Basal ganglia | 384 | -18 | -10 | 28 | -4.18908 | 79 |
| Basal ganglia | 560 | 14 | 28 | 0 | -4.40146 | 36 |
| Basal ganglia | 784 | 18 | -32 | 10 | -4.48621 | 44 |
| Cerebellum | 288 | 4 | -56 | -48 | -4.21364 | 100 |
| Cerebellum | 336 | 0 | -64 | -6 | -4.4739 | 57 |
| Cortex Default ModeA | 64 | -40 | -68 | 34 | -3.51907 | 38 |
| Cortex Default ModeA | 64 | -2 | -50 | 36 | -3.61961 | 50 |
| Cortex Default ModeA | 160 | -8 | 20 | -12 | -3.65508 | 50 |
| Cortex Default ModeA | 176 | -4 | 52 | -8 | -3.89427 | 68 |
| Cortex Default ModeA | 192 | 10 | -48 | 44 | -4.02499 | 46 |
| Cortex Default ModeB | 64 | -14 | 40 | 44 | -3.53953 | 50 |
| Cortex Default ModeB | 64 | -54 | -6 | 52 | -3.60081 | 88 |
| Cortex Default ModeB | 96 | -56 | 4 | -36 | -3.61481 | 100 |
| Cortex Default ModeB | 192 | -32 | 0 | -32 | -4.00529 | 21 |
| Cortex Default ModeB | 208 | -48 | 28 | 0 | -4.05867 | 81 |
| Cortex Default ModeB | 264 | -52 | -10 | 44 | -4.08027 | 48 |
| Cortex Dorsal AttentionA | 400 | 52 | -58 | 8 | -3.94604 | 88 |
| Cortex Dorsal AttentionA | 824 | 50 | -68 | -2 | -4.44292 | 24 |
| Cortex Dorsal AttentionB | 64 | -44 | -32 | 40 | -3.60794 | 75 |
| Cortex Dorsal AttentionB | 96 | 34 | -30 | 52 | -3.79994 | 58 |
| Cortex Dorsal AttentionB | 208 | -38 | 2 | 56 | -3.77187 | 88 |
| Cortex Dorsal AttentionB | 392 | 14 | -44 | 70 | -4.2037 | 49 |
| Cortex Dorsal AttentionB | 808 | 38 | 0 | 54 | -4.85951 | 63 |
| Cortex Fronto ParietalA | 128 | 32 | -48 | 52 | -3.75713 | 69 |
| Cortex Fronto ParietalA | 288 | 42 | 16 | 16 | -4.27468 | 8 |
| Cortex Fronto ParietalA | 336 | 38 | -2 | 40 | -4.4202 | 31 |
| Cortex Fronto ParietalA | 928 | 48 | 20 | 34 | -4.48172 | 31 |
| Cortex Fronto ParietalB | 128 | -6 | 20 | 40 | -3.84357 | 88 |

| | | | | | | |
|---|---|---|---|---|---|---|
| Cortex Fronto ParietalB | 176 | 8 | 44 | 36 | -4.37012 | 9 |
| Cortex Fronto ParietalB | 176 | 28 | 26 | 48 | -4.09105 | 82 |
| Cortex Fronto ParietalB | 200 | 58 | -56 | 46 | -3.92567 | 4 |
| Cortex Fronto ParietalB | 224 | 14 | 38 | -18 | -4.16718 | 18 |
| Cortex Limbic | 144 | -2 | 38 | -30 | -3.92036 | 44 |
| Cortex Limbic | 160 | 16 | 14 | -16 | -3.81417 | 30 |
| Cortex Limbic | 176 | 36 | 12 | -28 | -4.45452 | 77 |
| Cortex SomatomotorA | 64 | -62 | -10 | 22 | -3.55048 | 75 |
| Cortex SomatomotorA | 128 | -54 | -28 | 48 | -3.64743 | 50 |
| Cortex SomatomotorA | 904 | -30 | -20 | 60 | -4.93739 | 27 |
| Cortex SomatomotorB | 192 | 46 | -26 | 22 | -3.9968 | 71 |
| Cortex Temporal Parietal | 128 | -60 | -16 | -4 | -3.85981 | 31 |
| Cortex Ventral AttentionA | 64 | 34 | 14 | -18 | -3.68263 | 75 |
| Cortex Ventral AttentionA | 112 | -54 | -36 | 48 | -3.72868 | 57 |
| Cortex Ventral AttentionA | 128 | 12 | -12 | 44 | -3.78576 | 44 |
| Cortex Ventral AttentionA | 192 | 66 | -20 | 40 | -3.74956 | 8 |
| Cortex Ventral AttentionA | 192 | -2 | 0 | 56 | -3.7675 | 46 |
| Cortex Ventral AttentionB | 64 | -20 | 44 | -18 | -3.56508 | 100 |
| Cortex Ventral AttentionB | 64 | 36 | 22 | 8 | -3.59033 | 100 |
| Cortex Ventral AttentionB | 616 | 4 | 36 | 30 | -4.72523 | 60 |
| Cortex Visual Central | 192 | 42 | -80 | -4 | -3.94547 | 88 |
| Cortex Visual Central | 336 | 0 | -84 | -12 | -3.86713 | 48 |
| Cortex Visual Central | 344 | -42 | -76 | 2 | -4.60834 | 40 |
| Cortex Visual Peripheral | 344 | 6 | -40 | -2 | -4.07015 | 37 |
| Diencephalon | 176 | -2 | -4 | 6 | -3.5862 | 36 |
| Diencephalon | 272 | 0 | -26 | 8 | -4.42874 | 18 |
| Hippocampus | 96 | 12 | -38 | 4 | -3.57135 | 17 |
| No description | 64 | -44 | -46 | -4 | -3.605 | 0 |
| No description | 64 | 16 | 46 | -4 | -3.64018 | 0 |
| No description | 64 | -30 | 16 | 32 | -3.61792 | 0 |
| No description | 128 | -20 | -44 | 16 | -3.79467 | 0 |
| No description | 224 | -26 | -46 | 12 | -4.45522 | 0 |

*Note.* The "% covered" column indicates the percent of voxels in a cluster cover by a specific region label. Automated labelling was performed with the canlab toolbox using the region() and table() functions.

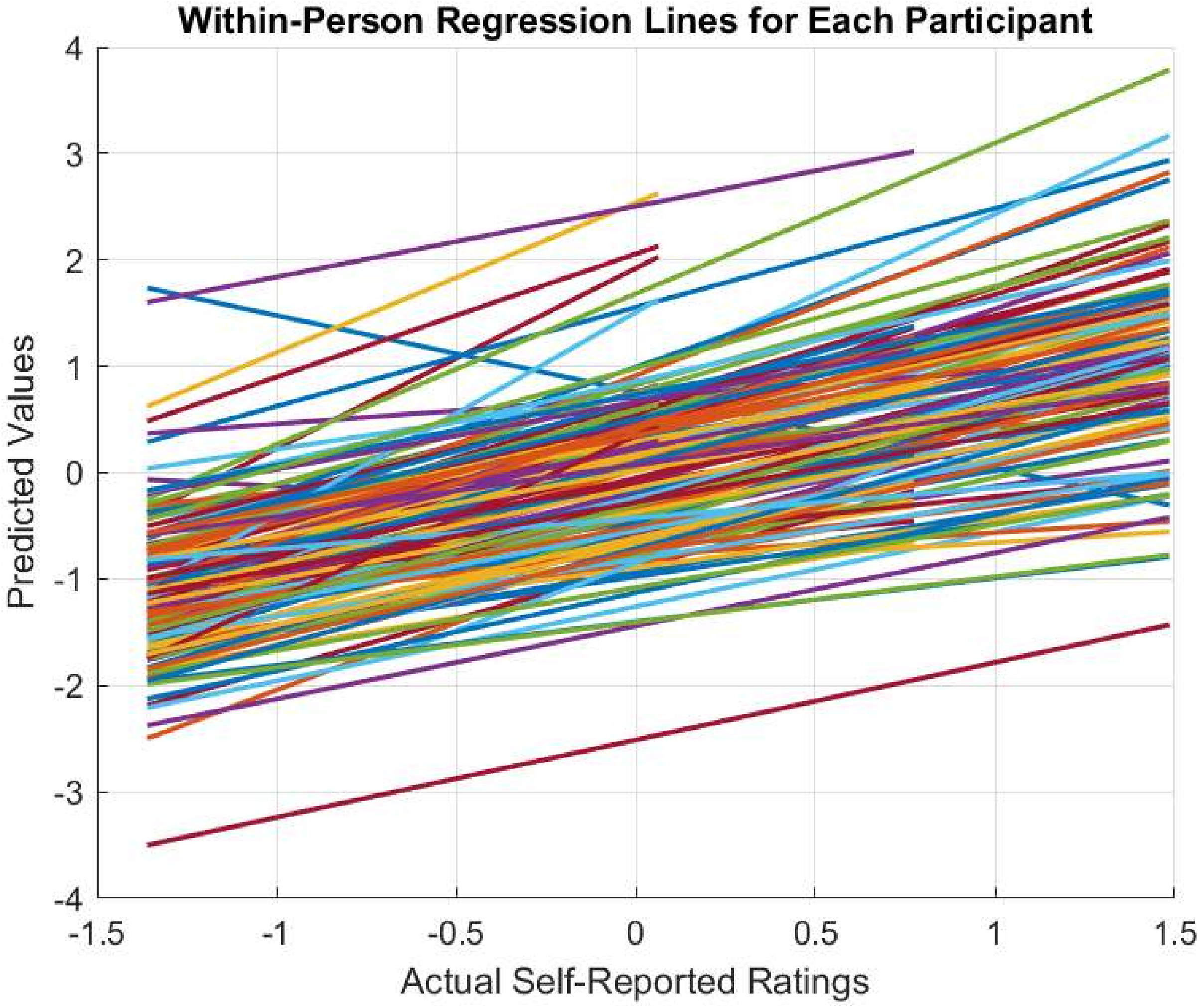

**Figure S1.**
Within-person regression lines for the association between actual state-based self-report rating of negative affect and predicted values from a support vector regression.
